# Micro-community associated with ectomycorrhizal *Russula* symbiosis and sporocarp-producing *Russula* in Fagaceae dominant nature areas in southern China

**DOI:** 10.1101/2020.04.22.056713

**Authors:** Wen Ying Yu, Ming Hui Peng, Jia Jia Wang, Wen Yu Ye, Zong Hua Wang, Guo Dong Lu, Jian Dong Bao

## Abstract

*Russula griseocarnosa*, an ectomycorrhizal (ECM) fungus, is a species of precious wild edible mushrooms with very high market value in southern China. Its yield is affected by many factors including the tree species and environmental conditions such as soil microbiome, humidity. How the microbiome promotes the ECM fungus symbiosis with Fagaceae plants and sporocarp-producing has never been studied. In this study, we collected rhizosphere samples from Fujian province, the microbiota in the root and mycorrhizal rhizosphere were identified by Illumina MiSeq high-throughput sequencing. First, we compared three types of fungal communities: root tips infected with ECM *Russula* (type 1), tips with *Russula* sporocarp (type 2) and tips without ECM (type 3). Our results showed that the fungal richness was negatively correlated with *Russula. Russula, Tomentella* and *Lactarius* were common in Fagaceae ECM roots. As to the mycorrhizal interactions, *Boletu*s may be considered as an indicator species for sporocarp-producing *Russula*, and *Acremonium, Cladophialophora* were associated with *Russula* symbiosis. Second, we analyzed the fungal and bacterial communities in rhizosphere soils from the corresponding to previously three types (type 1, 2, 3). *Dacryobolus* and *Acidocella* may be considered as an indicator species for sporocarp-producing *Russula*. Fungi *Tomentella, Saitozyma, Elaphomyces* and bacteria *Acidicaldus, Bryobacter, Sorangium* and *Acidobacterium* occurred more frequently in the ECM *Russula* rhizosphere. Furthermore, the indicators *Elaphomyces, Tomentella, Sorangium* had a positive correlation with *Russula* symbiosis by network analyses. Overall, our results suggest a relationship between micro-community and ECM *Russula* formation and *Russula* sporocarp, which may provide new strategies for improving *Russula* symbiosis rate and sporocarp production.

## IMPORTANCE

*Russula* (Russulaceae, Russulales, Agaricomycetes, Basidiomycota) species are ectomycorrhizal (ECM) fungi that form symbiotic associations with host roots. Approximately 750 *Russula* species have been reported in worldwide (1). *Russula* taxa showed high diversity, strong habitat preference and some preference for soil horizons (2)(Jo’zsef Geml 2010). Li M(3)(2010)identified that *Russula* emerge at least three divergent lineages based on the genetic diversity and geographic differentiation in southern China, and *R. griseocarnosa* belongs to one of the lineages. *R. griseocarnosa* is described from southern China, including Guangdong, Yunnan, and Fujian Provinces (4). *R. griseocarnosa* sporocarp, which is popularly commercially as food and medicine, is uncultivatable and collected from the natural habitat. To date, there is little knowledge about the controlled production of *Russula* and its micro-community.

Several biotic factors affect ectomycorrhizal fungal communities: plant host species (5, 6), plantation age (7), asymptomatic ectomycorrhiza endophytes (8), ectomycorrhizal propagules in the soil(9) and the traits of the dispersal propagules of ECM fungi (10). The common ectomycorrhizal *Russula* sp. was associated with seven host genera: *Pleioblastus chino, Quercus serrata, Symplocos prunifolia, Ilex pedunculosa, Prunus jamasakura, Gamblea innovans*, and *Lyonia ovalifolia*(11, 12). *R. griseocarnosa* grows in forests with Fagaceae in southern China (13). ECM fungal communities strongly vary during long-term ecosystem development, even within the same hosts (14). Some *Russula* spp. was reported to be dominant colonizers of mature roots of pines (15)(Horton & Bruns, 2001) but did not colonize bioassay seedlings (16).’Early stage’ and ‘late stage’ mycorrhizal fungi of birch appeared distinction (7). Gebhardt also found that Ectomycorrhiza communities of *Quercus rubra* of different age performed specificity with low similarity between the chronosequence stands site (17). Some fungal root endophytes prefer ECM formed by particular species of ectomycorrhizal fungi (8, 18), potentially representing a second level of root endophyte selection. Dark septate endophytes are considered to be ubiquitous, colonizing mycorrhizal(19). ECM propagule communities in soil may diverge from those root-colonizing ECM communities and affect persistence of symbiotic relationship between mycorrhiza and available host roots by competitive networks(20). ECM propagules in soil are less frequent and diverse in early primary succession and become more frequent and diverse along forest development, due mainly to the accumulation of dormant spores of *Rhizopogon* spp. and sclerotia of *Cenococcum* spp(20). Simultaneously dispersal ability across ECM species correlated well with the composition of communities associated with host (21). *Russula* naturally propagate by short-distance spore dispersal rather than vegetative growth of dikaryophytic mycelia or long-distance spore dispersal(22). In all, the *Russula* ECM community may be affected by symbiotic structures and root endophytes in the roots, mycorrhizal extraradical mycelia, spores in the soil, as well as its dispersal ability.

Mycorrhizal interactions are usually classified on the basis of the features of the symbiotic interfaces(23). The ECM community colonizing root tips was strongly structured by competitive interactions or ecological processes generating a similar spatial pattern, rather than neutral processes. The ECM *Cortinarius* sp. and *Lactarius rufus* competed for root tips(24). Both *Cenococcum geophilum* and *Clavulina cinerea* as mycorrhizas and as extramatrical mycelium (EMM) in a *Pinus resinosa* plantation showed a negative correlation(25). Hortal suggests that interspecific competition between *Lactarius deliciosus* (inoculated fungi) and *Rhizopogon roseolus* (indigenous fungi) occurs in the root system for ectomycorrhiza formation in available roots, rather than in the extramatrical mycelium phase (26). However, associations among *Suillus bovinus* and *Gomphidius roseus* occur within ectomycorrhizal roots (27). There are also a number of notable examples of associations between sporocarps of different species, such as between *Boletus parasiticus* and *S. citrinum, Asterophora parasitica* and *Lactarius*, and between *Claudopus parasiticus* and *Cantharellus cibarius* (28),. However, mycorrhizal interaction studies of *Russula* are limited in the literature.

Similar to the rhizosphere, mycorrhizosphere is the feet of fruiting bodies or the root of ectomycorrhizal fungi. Mycorrhizosphere may constitute a hot spot(29), in which microorganisms were affected(30-32).The mycosphere effect was prominent, though previous studies have failed to demonstrate significant differences between bacterial communities associated with roots colonized by *Suillus variegatus* and *T. submollis, Lactarius* sp. and *Tomentella* sp.(33) or by *C. geophilum* and *Russula* sp. (34). It has also been established that mycorrhizal networks have different microbial communities compared with bulk soil by stimulating some families such as *Bradyrhizobiaceae, Burkholderiaceae*, and *Pseudomonadaceae* (35). Bacterial activity is stimulated by the provision of easily available nutrition such as carbonaceous compounds (30).The mycosphere effect might exert strong influences on the bacterial community of soil(36), in which bacterial activity is stimulated by trehalose, which is degraded by hyphosphere and derived from fungi (37, 38). Andersson(39)(2003)., who used phospholipid fatty acid profiles to characterize bacterial communities, has demonstrated that basidiomycete wood-decomposing fungi are able to influence bacterial community structure. Bacteria adapted to the mycospheres of three or more or just one fungal species was defined as specific selective bacterial(40). The specific members of the Sphingomonadaceae family selected were at the bases of the fruiting bodies of the ectomycorrhizal fungi *Laccaria proxima* and *Russula exalbicans* in comparison to the adjacent bulk soil, major of which did not cluster with known bacteria from the database (36, 41). On the other hand, some mycorrhiza-associated bacteria have been shown to produce compounds that are antagonistic to plant pathogens (42). Meanwhile, “mycorrhiza helper bacteria” (MHB) appeared to positively influence the development and function of ectomycorrhiza (43), but the effects varied with different species combinations (44, 45). *Ralstonia* sp. and *Bacillus subtilis* can promote *Suillus granulatus*-infected *Pinus thunbergii* (34). *Bacillus subtilis* helped the growth of *Cenococcum geophilum* Fr and promoted *Suillus granulatus* infection (34) but inhibited *Rhizopogon* sp. infection (45). Apart from bacterial, ectomycorrhizal symbionts strong also selected ascomycete communities and other ECMs(31) (32).Högberg (2002)(46) demonstrated that the EMM is at least 30% of the microbial biomass in boreal forest soils. ECM fungi competed with saprotrophic fungi in soil by the EMM(47). *Tuber rufum* and some members of *Boletales* are typically restricted to productive truffle plots. On the other hand, *Hebeloma, Laccaria* and *Russula* species are mostly associated with unproductive truffle plots, Ectomycorrhizae belonging to *Thelephoraceae* are frequently found in mature truffle orchards but do not seem to affect sporocarp production (32). Ascomycetes associated with ectomycorrhizas: molecular diversity and ecology with particular reference to the Helotiales had been reported (48).However, the rhizosphere, mycosphere effect of *Russula griseocarnosa* has not been recorded in the literature.

Regarding the relationship between the amount of soil EMM with ectomycorrhizae and sporocarps, different mycorrhizas have different conclusions. Zampieri et al.(2010)(49) showed that the mycelium of *Tuber magnatum* was more widespread than was inferred from the distribution of its fruiting bodies and ectomycorrhizae. Zhou et al. (2001) (50)demonstrated that the development of *Suillus grevillei* sporocarps is correlated with amount of EMM and ectomycorrhizae of *S. grevillei* in a narrow area. The *Tuber melanosporum* EMM biomass detected in the soil from the natural truffle ground was significantly greater than that of other plant orchards analyzed, and the lowest amount of *T. melanosporum* mycelium maintained a sporocarp production in plant orchards (51). However, Some ECMs are consistent in ectomycorrhizae but inconsistent in sporocarps. De la Varga et al. (52)(2012) quantified *B. edulis* extraradical mycelium in a Scots pine forest and found a positive correlation between the amount of mycelia and the presence of *Boletus edulis* mycorrhizae, but not with the productivity of fruiting bodies. However, the relationship between the amount of *Russula* with ectomycorrhizae and sporocarps is scarce in the literature.

The relation between the productivity of fruiting bodies and ECM symbiosis is still unclear. Some ectomycorrhizal species produce abundant ectomycorrhizal root tips but few or no fruiting bodies, while other ectomycorrhizal fungi form abundant fruiting bodies but a low number of ectomycorrhizal root tips. Guidot et al. (53)(2001) found a spatial congruence of above- and belowground distribution for *H. cylindrosporum*. However, De la Varga et al.(52) (2012) found that the presence of mycorrhizae of the *B. edulis* symbiotic rate was not consistent with the production of fruiting bodies. There are also *Russula* species difference between the above-ground *Russula* sporocarp and underground *Russula* mycorrhizal(2). Geml(2010) (2)observed that 15 and 45 of the 50 *Russula* phylogroups species were found in sporocarp and soil clone sequences, respectively. Given the long delay between the establishment of the plantation and the formation of sporocarps, short- and medium-term control of the survival and persistence of fungal symbionts in plantations have to still be evaluated by the assessment of vegetative structures as the ectomycorrhizas or extraradical mycelium.

Microbial community affected mycorrhizal fungal function (for example, symbiosis establishment capacity, sporocarp production), and the reciprocal effects are vice versa. The microorganisms associated with mycorrhizal fungi may either have positive or negative impacts on fungal spore germination, growth, nutrient acquisition and plant colonization (43, 54). *Tuber indicum* altered the ectomycorrhizosphere and endoectomycosphere microbiome and metabolic profiles of the host tree *Quercus aliena* (55). *Tuber borchii* shapes the ectomycorrhizosphere microbial communities of *Corylus avellana*(56). Therefore, we speculate that detecting the communities of *Russula* can tracked ECM persistence throughout the entire biological cycle, which will help to control ectomycorrhiza formation and sporocarp production. In this study, to understand the communities of the targeted *Russula griseocarnosa* and to find the possible indicator microbes of successful *Russula griseocarnosa* plantations, we identified the *Russula* ectomycorrhizal fungal communities inhabiting different life cycle stages based on MiSeq sequencing of ribosomal internal transcribed spacer (ITS) sequences of root DNA and mycosphere communities based on MiSeq sequencing of the 16S V3-V4 as well as ribosomal internal transcribed spacer (ITS) sequences of mycosphere soil DNA.

This study is the first attempt to analyze ectomycorrhizal communities of *Russula* using MiSeq sequencing metagenomics DNA of *Russula* root and the Mycorrhizal rhizosphere soil in *Russula* at different stages. We think that *Russula* shapes the ectomycorrhizosphere microbial communities of Fagaceae *(Quercus glauca* and *Castanopsis hainanensis*.).

## RESULTS

### Comparing microbiomes among types of the ECM fungus Russula

ECM *Russula* was first identified by combining morph typing with Sanger sequencing DNA sequences. We analyzed internal transcribed spacer (ITS) rDNA sequences of ECM root tips and ECM rhizosphere soil samples using phylogenetic methods, operational taxonomic unit (OTU) delimitations and ordinations to compare species composition in various types of ECM *Russula*.

In the ECM metagenome, we found a positive correlation between the concentration of *Russula* DNA and the presence of *Russula* mycorrhizae in the mycorrhizal rhizosphere and ECM root (Tab 1). To analyze whether distinctive communities are selected by *Russula* ectomycorrhizal fungi, the fungi of ECM root microbiomes were compared and divided into three types. *Russula* that could be detected by Sanger sequencing with a DNA concentration above 10% of total ECM genomic DNA, as determined by MiSeq sequencing, were classified as type 1. *Russula* sporocarps that could be collected from the ground and the extended hyphae of which could be connected between roots of the host and sporocarp within 50 cm, with *Russula* detected by Sanger sequencing, were classified as type 2. *Russula* that were not detected by Sanger sequencing or with a DNA concentration below 5% of the total ECM genomic DNA, as determined by MiSeq sequencing, were classified as type 3. Therefore, the samples of ECM tip and rhizosphere samplings were divided into three types: *Russula*-infected (type 1), sporocarp-producing Russula (type 2), and Russula-uninfected (type 3) (Tab 1). Relative *Russula* OUT abundance is significant difference in ECM *Russula* symbiosis root and in ECM *Russula* rhizosphere, respectively (Tab 1, Fig. 1A).Type 2 is the most abundant, type 1 is the second, and type 3 is the least. Interestingly, in type 3, the amount of *Russula* in the soil is higher than in the root. This may indicate that in the natural growth area of *Russula*, there are a large number of *Russula* propagules in the soil.

**TABLE 1.**
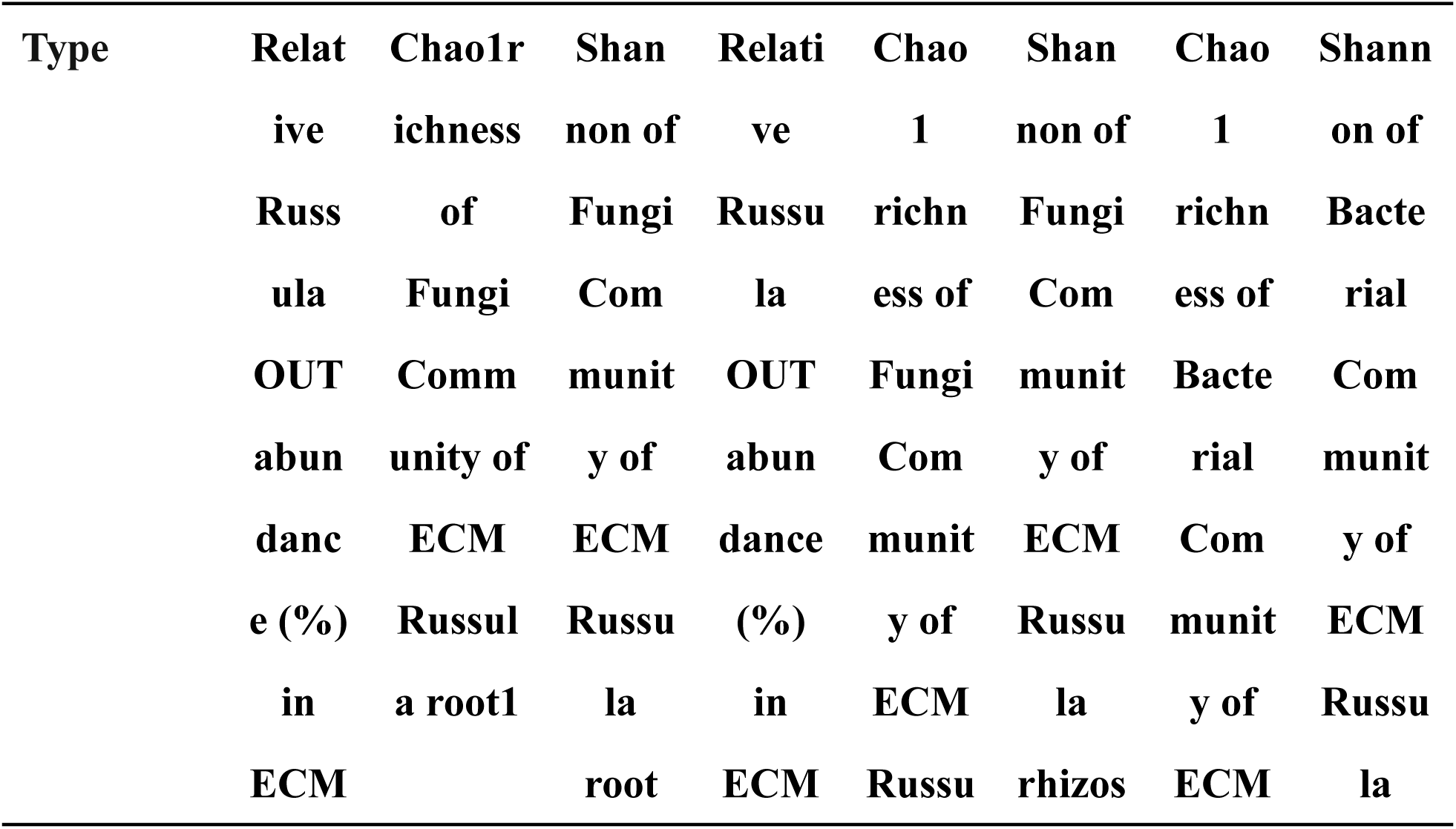
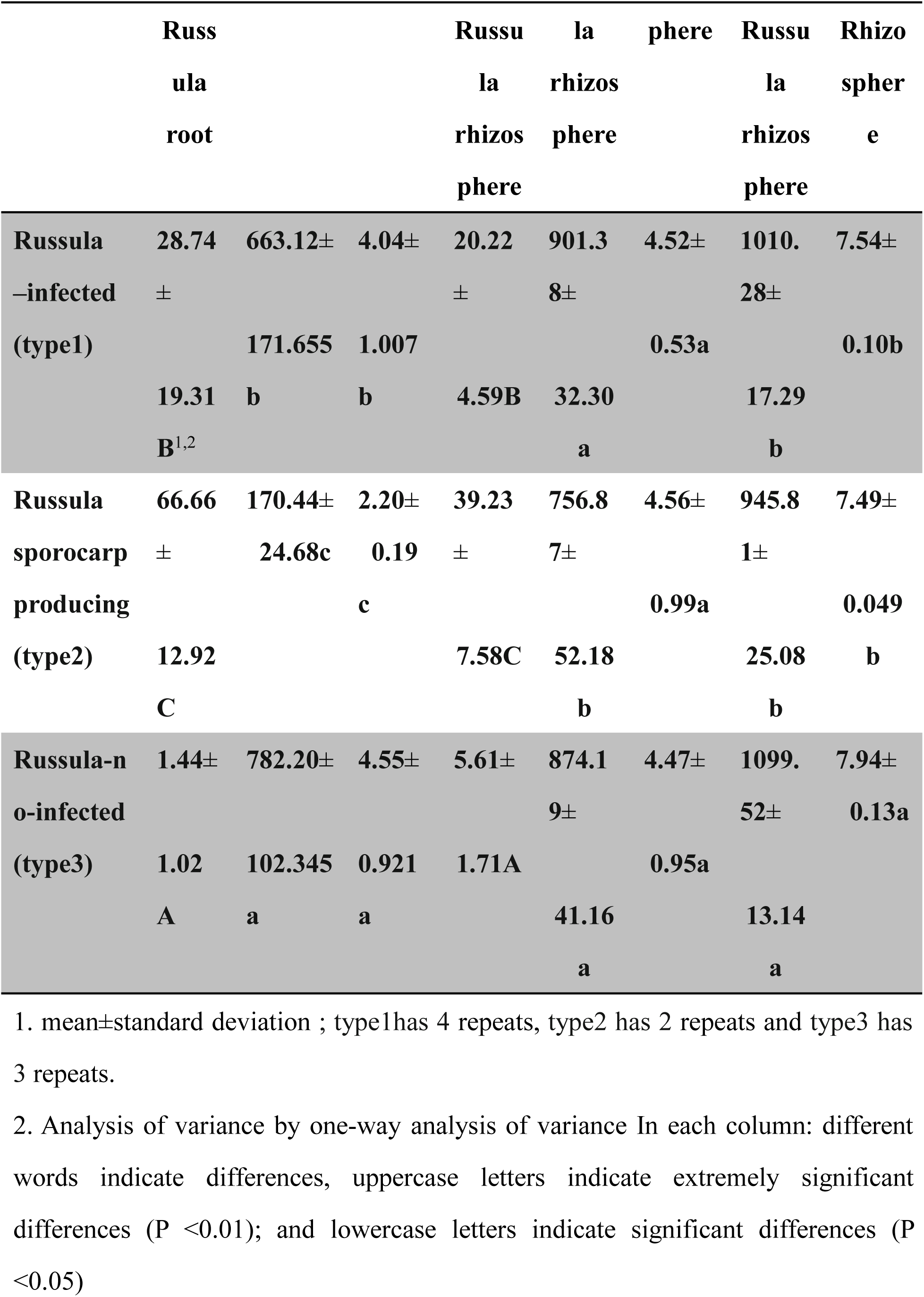
Relative *Russula* OUT abundance in three types and Alpha diversity analysis of fungi and bacterial of three types of ECM *Russula* roots or *Russula* rhizosphere soil in natural *Russula* growth areas

| Type | Relative<br>Russula<br>OUT<br>abundance<br>(%)<br>in<br>ECM | Chao1<br>richness<br>of<br>Fungi<br>Community<br>of<br>ECM<br>Russula<br>root | Shannon<br>of<br>Fungi<br>Community<br>of<br>ECM<br>Russula<br>root | Relative<br>Russula<br>OUT<br>abundance<br>(%)<br>in<br>ECM | Chao1<br>richness<br>of<br>Fungi<br>Community<br>of<br>ECM<br>Russula<br>rhizosphere | Shannon<br>of<br>Fungi<br>Community<br>of<br>ECM<br>Russula<br>rhizosphere | Chao1<br>richness<br>of<br>Bacterial<br>Community<br>of<br>ECM<br>Russula<br>rhizosphere | Shannon<br>of<br>Bacterial<br>Community<br>of<br>ECM<br>Russula<br>rhizosphere |
| --- | --- | --- | --- | --- | --- | --- | --- | --- |

|  | Russ<br>ula<br>root |  |  | Russu<br>la<br>rhizos<br>phere |  |  | Russu<br>la<br>rhizos<br>phere |  |
| --- | --- | --- | --- | --- | --- | --- | --- | --- |
| **Russula** | **28.74** | **663.12±** | **4.04±** | **20.22** | **901.3** | **4.52±** | **1010.** | **7.54±** |
| **–infected** | **±** |  |  | **±** | **8±** |  | **28±** |  |
| **(type1)** |  | **171.655** | **1.007** |  |  | **0.53a** |  | **0.10b** |
|  | **19.31** | **b** | **b** | **4.59B** | **32.30** |  | **17.29** |  |
|  | **B<sup>1,2</sup>** |  |  |  | **a** |  | **b** |  |
| **Russula** | **66.66** | **170.44±** | **2.20±** | **39.23** | **756.8** | **4.56±** | **945.8** | **7.49±** |
| **sporocarp** | **±** | **24.68c** | **0.19** | **±** | **7±** |  | **1±** |  |
| **producing** |  |  | **c** |  |  | **0.99a** |  | **0.049** |
| **(type2)** | **12.92** |  |  | **7.58C** | **52.18** |  | **25.08** | **b** |
|  | **C** |  |  |  | **b** |  | **b** |  |
| **Russula-n** | **1.44±** | **782.20±** | **4.55±** | **5.61±** | **874.1** | **4.47±** | **1099.** | **7.94±** |
| **o-infected** |  |  |  |  | **9±** |  | **52±** | **0.13a** |
| **(type3)** | **1.02** | **102.345** | **0.921** | **1.71A** |  | **0.95a** |  |  |
|  | **A** | **a** | **a** |  | **41.16** |  | **13.14** |  |
|  |  |  |  |  | **a** |  | **a** |  |

1. mean±standard deviation ; type1has 4 repeats, type2 has 2 repeats and type3 has 3 repeats.
2. Analysis of variance by one-way analysis of variance In each column: different words indicate differences, uppercase letters indicate extremely significant differences (P <0.01); and lowercase letters indicate significant differences (P <0.05)

**Fig. 1.**
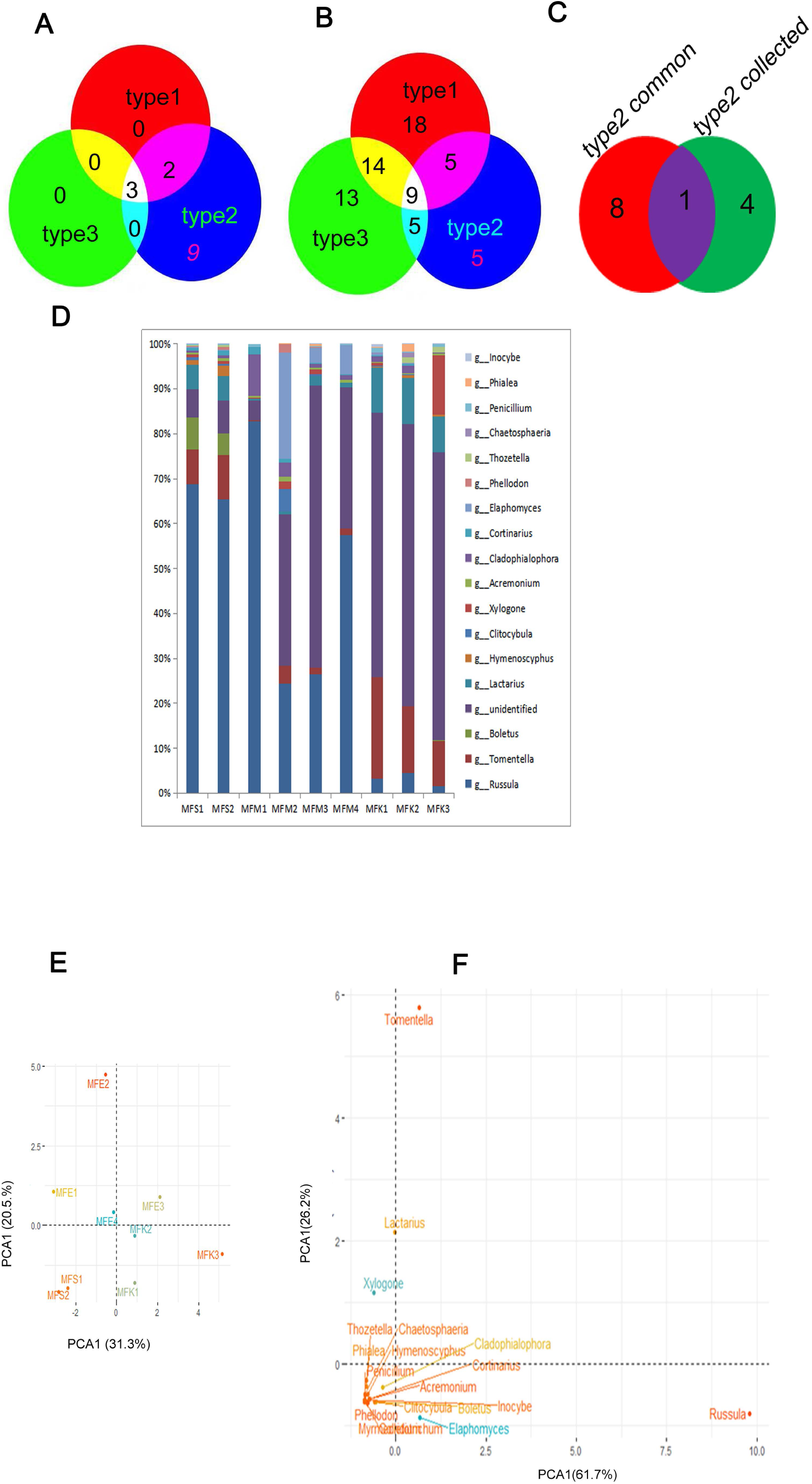
Diversity analysis of fungi of three types of *Russula* symbiosis roots in natural Russula growth areas. A:The main fungi genus composition and abundance in type2 *Russula* symbiosis root and the corresponding genus are in the other two types B: The Venny mapping of the common genus of the top 20 fungi genus in three types Russula symbiosis root C: The Venny mapping of the collection genus of the top 20 fungi genus in three types Russula symbiosis root D: The Venny mapping of the common genus and the collection genus of the top 20 fungi genus in type2 Russula symbiosis root E : Analysis of ECM role difference of three types Russula symbiosis root on top 20 fungi genus of ECM Russula by PCoA F : Analysis of composition difference of three types Russula symbiosis root on top 20 fungi genus of ECM Russula by PCoA In Eand F: The scales of the horizontal and vertical axes are relative distances and have no practical significance. The contribution rate is the degree of interpretation, and the hypothetical factors can be evaluated and verified. In A,Eand F :In the sample name, the first letter M indicates mycorrhizal root; first second F stands for fungi; the third letter different treatment :S indicates sporocarp(type2), E indicates ECM ectomycorrhizal(type1), K indicates control(type3) respectively; the fourth number indicates the different biological repetitions of different treatment.

### Fungal diversity analysis and analysis of indicator species associated with ECM *Russula* roots

In total, 1346 fungal operational taxonomic units (OTUs) were distinguished in roots. Among the three types, the Chao1 diversity index and Shannon diversity index decreased in sporocarp-producing *Russula* type (type 2) and *Russula*-infected type (type 1) compared to the *Russula*-uninfected (type 3) (Tab 1). The result shows that species richness shifted in the composition of the ECM community associated with *Russula*.The fungal community composition and the abundance of the main fungi (over 0.05% fungi in roots genome) were different in the three *Russula* types roots (Fig. 1A,Tab 1).The result shows that *Russula* is dominates in fruiting bodies (type 2) and infected samples(type 1) though *Russula* in three type samples.

Based on the top 20 fungi of each sample, we first obtained the common species of three types after intersecting each type with the Venny mapping tool: 14 species were in type 2, 5 in type 2 and 3 in type 3 (Tab S1). Then, we intersected the common types of three types with the Venny mapping tool. At the genus level, the three types shared 3 common genera: *Russula, Tomentella* and *Lactarius* (Tab S2, Fig. 1B).These results showed that these are common fungi in Fagaceae ECM roots. Types 1 and 2 shared 2 common genera: *Acremonium* and *Cladophialophora*, in addition to the abovementioned three common ECM(Tab S2,Fig. 1B). Second, We take a collection of three types genera respectively, subsequently we intersected the collection of three types by the Venny mapping tool (Fig. 1C): 24 species in type 2, 46 in type 1 and 41 in type 3, and 5 genera were exclusively in type 2 collection. We take the intersection of the unique collection (9 genera) and union (5 genera) in type 2, and found only one species boletus (Fig. 1D, Tab. 2). Analyze fungi composition differences of three types roots by PCoA based on the top 20 fungal genera, we found that the control (type 3) belonged to quadrant IV, type 1 belonged to quadrant I or II, and type 2 belonged to quadrant III (Fig. 1F). The results show that the fungal community of sporocarp-producing *Russula* was completely different from that of Russula-uninfected. The *Russula*-infected type was in the transitional phase.We suggest the *Russula* infection contributed 31.3% and others elements host contributed 20.5% of the differences, respectively. *Boletus* was the only ECM with the emergence of *Russula* fruit bodies. In all, *Boletus* may be considered an indicator species in the *Russula* sporocarp-producing fungal community, and *Acremonium* and *Cladophialophora* may be considered indicator species in *Russula* symbiosis fungal communities (Tab. 2). Analyze the different effects of ECM species by PCoA based on the top 20 fungal genera, we found *Tomentella, Xylogone*, and *Lactarius* belonged to I, II, while others belonged to quadrant III,V (Fig. 1E). The *Russula* infection contributed 55% and others elements contributed 24% of the differences, respectively. Combining these two factors, it shows *Russula* and *Elaphomyces* can be divided into one categories, while *Xylogone* can be divided into another category functionally. So we assume that *Elaphomyces* are benefit for *Russula* symbiosis while *Tomentella, Xylogone*, and *Lactarius* has the function of competing hosts.

**Table 2.**
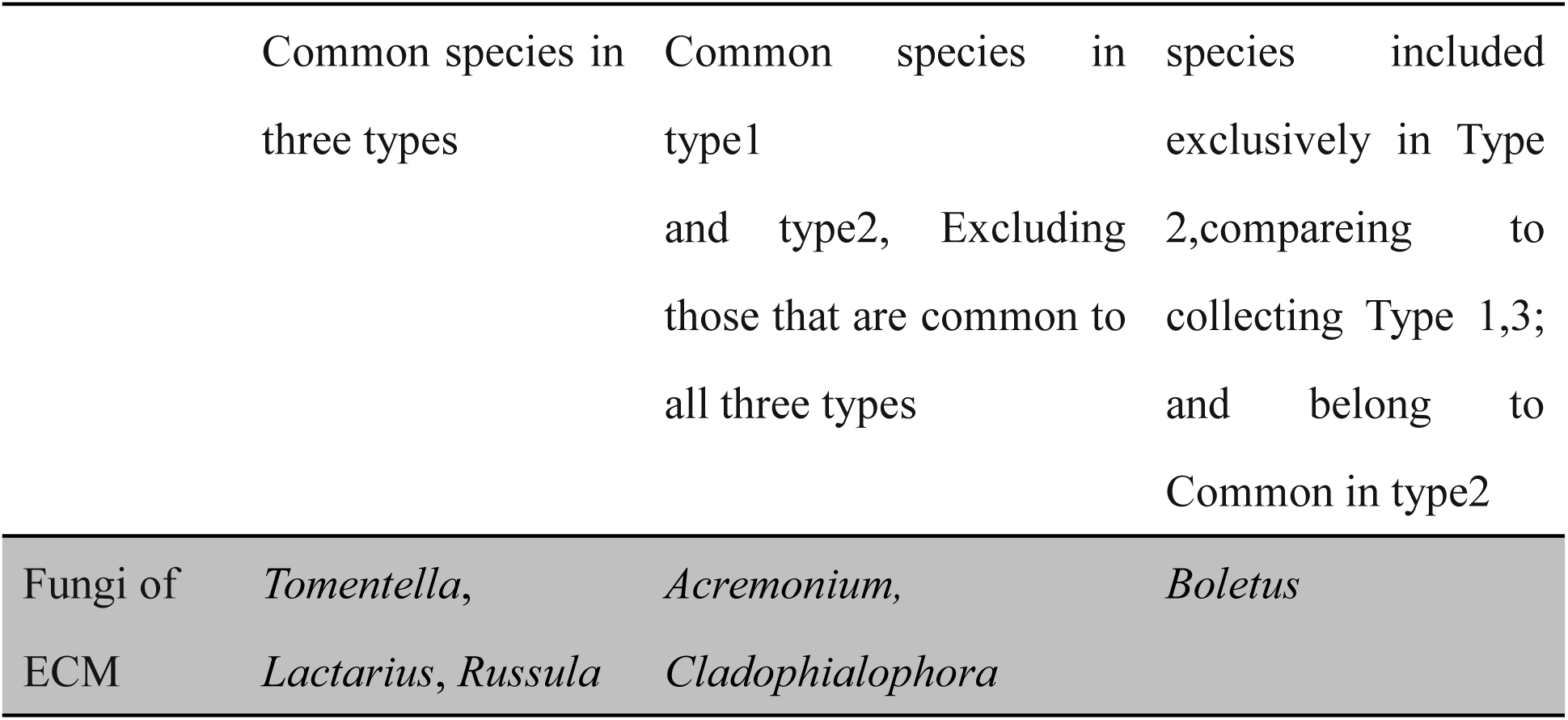
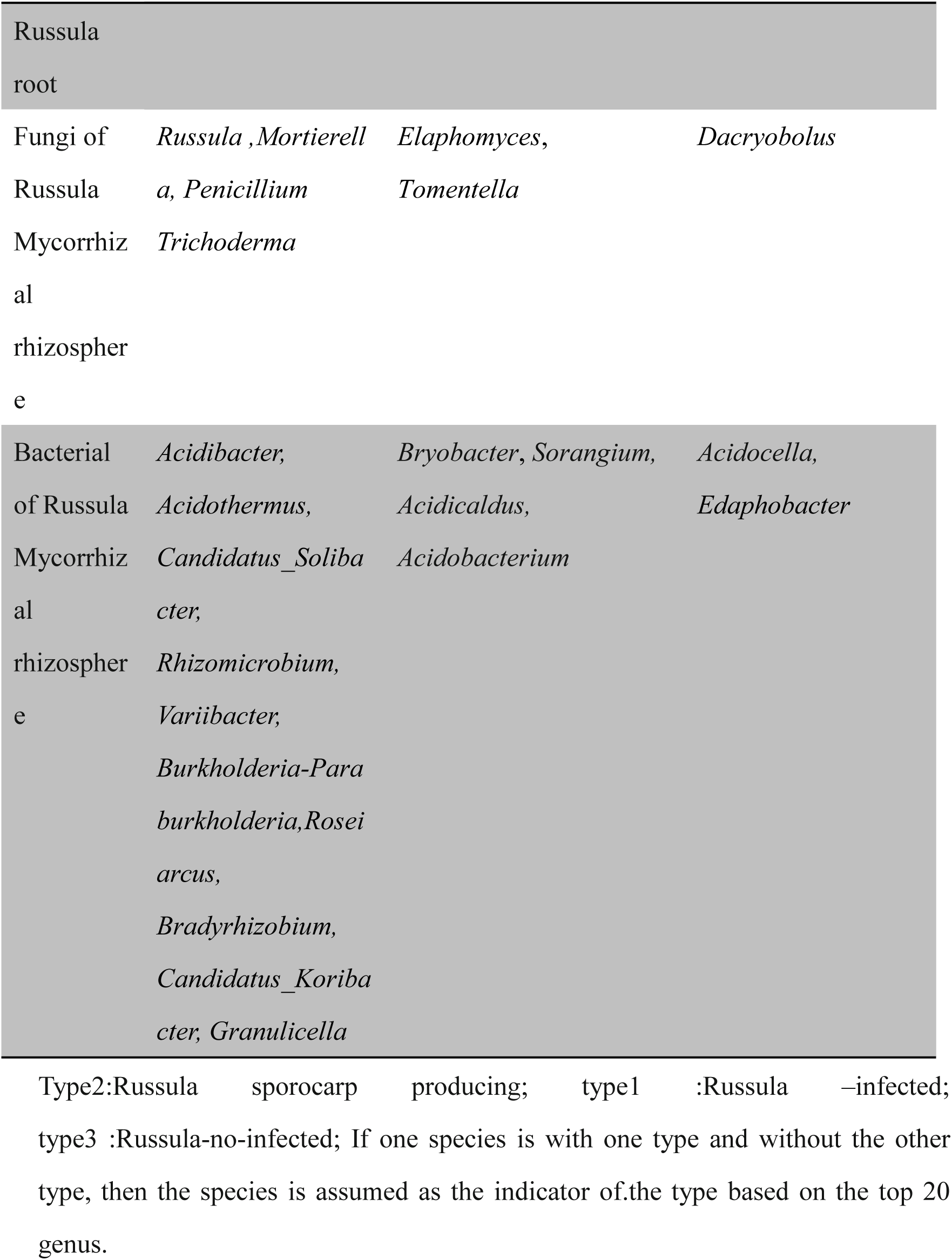
Indicator species of fungi and bacterial community of Russula Mycorrhizal based on the top 20 genus

### Fungal diversity analysis and analysis of indicator species associated with ECM *Russula* rhizosphere soil

In total, 1829 fungal operational taxonomic units (OTUs) were distinguished in rhizosphere soil. Interactions with native ectomycorrhizal fungi present in the soil play a key role in the higher diversity of fungal taxa .Compared to type 1 and 3, the Chao1 diversity index decreased in types 2; while there were no significant differences between types 1 and 3 (Tab. 1). But, the Shannon diversity index showed no significant differences in three types (Tab. 1). The fungal community composition and the abundance of the main fungi (over 0.05% fungi in soil genome)were different in the three *Russula* types rhizosphere soil (Fig. 1A,Tab 1).Compared to the root, The result shows that *Russula* is also dominates in fruiting bodies (type 2) and infected samples(type 1) though *Russula* in three type samples(Fig. 1A,Fig. 2A). However,compared to the root, the genera variety increased (Fig. 1A,Fig. 2A).

**Fig. 2.**
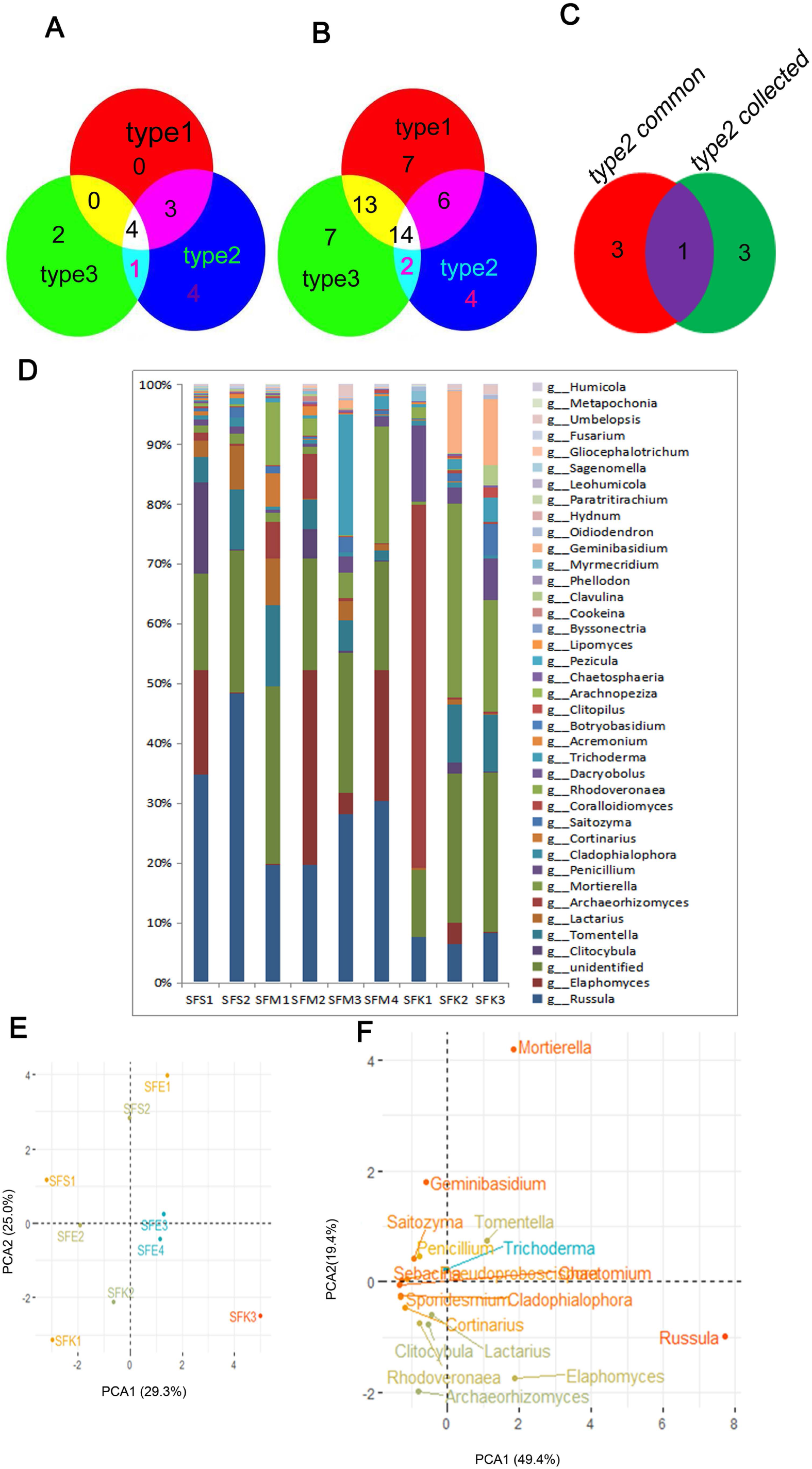
Diversity analysis of fungi of three types *Russula* mycorrhizal rhizosphere in natural *Russula* growth areas. A:The top 20 fungi genus **c**omposition and abundance in type2 *Russula* symbiosis mycorrhizal rhizosphere and the corresponding genus are in the other two types in at Fujian,China In the sample name, the first letter S indicates mycorrhizal root soil; first second F stands for fungi; the third letter different treatment :S indicates sporocarp, E indicates ECM ectomycorrhizal, K indicates control respectively; the fourth number indicates the different biological repetitions of different treatment. B: The Venny mapping of the common genus of the top 20 fungi genus in three types *Russula* symbiosis Mycorrhizal rhizosphere C: The Venny mapping of the collection genus of the top 20 fungi genus in three types *Russula* symbiosis Mycorrhizal rhizosphere D: The Venny mapping of the common genus and the collection genus of the top 20 fungi genus in type2 *Russula* symbiosis Mycorrhizal rhizosphere E : Analysis of ECM role difference of three types Russula symbiosis mycorrhizal rhizosphere on top 20 fungi genus of ECM Russula by PCoA F :Analysis of composition difference of three types Russula symbiosis mycorrhizal rhizosphere on top 20 fungi genus of ECM Russula by PCoA In E and F: the scales of the horizontal and vertical axes are relative distances and have no practical significance. The contribution rate is the degree of interpretation, and the hypothetical factors can be evaluated and verified.

The community composition of the top 20 fungi in the three *Russula* rhizosphere soil types was analyzed (Tab. S2). We first obtained the common species of three types after intersecting each type using the Venny mapping tool: 12 species in type 2, 7 species in type 1 and 7 species in type 3 (Fig. 2B). Then, we intersected the common types of the three types with the Venny mapping tool. At the genus level, the three types shared 4 common genera: *Russula, Mortierella, Penicillium* and *Trichoderma*; and *Russula* also had a large frequency in types 1 and 2 (Fig. 2B). This showed that 4 genera are common fungi in Fagaceae-dominant rhizosphere soil. Types 1 and 2 shared 3 common genera: *Elaphomyces, Tomentella* and *Saitozyma*, in addition to the abovementioned 4 common genera(Fig. 2B). Second, we take a collection of three types genera respectively, subsequently intersected the collection of three types by the Venny mapping tool (Fig. 2C): 26 species in type 2, 40 in type 1 and 36 in type 3; 4 genera were included exclusively in the type 2 collection. We take the intersection of the unique collection (4 genera) and union (4 genera) in type 2, and found *Dacryobolus* unique belonged to type 2 (Fig. 2D, Tab. 2). The top 20 fungal genera in the sporocarp-producing *Russula* root rhizosphere with types 3 or types 1 were compared. The result shows that *Russula* EMM is dominates in fruiting bodies (type 2) and infected samples (type 1) though *Russula* in three type samples. *Elaphomyces* also dominated in types 1 and 2, but nearly did not exist in the control, type 3 (Fig. 2D). In all, *Dacryobolus* may be considered an indicator species of sporocarp-producing *Russula* in the *Russula* rhizosphere. *Elaphomyces*, and*Tomentella* may be considered indicator species for *Russula* symbiosis in the rhizosphere (Tab. 2). Analyze three types of fungi composition differences of top 20 genera by PCoA in *Russula* rhizosphere soil based on the top 20 fungal genera, we found that type 2 belonged to quadrant II and type 1 belonged to quadrants I, III and IV, while type 3 belonged to quadrants III and IV, down the horizontal axis (Fig. 2E). The host and *Russula-*infection contributed 29.3% and 25.0%, respectively. Furthermore, analyzing the different effects of fungi species by PCoA, we found *Russula* and *Elaphomyces* belonged to quadrant IV, *Mortierella* and *Tomentella* belonged to quadrant I, *Trichoderma, Penicillium, Geminibasidium* and *Saitozyma* belonged to quadrant II, while others belonged to quadrant III (Fig. 2E). The *Russula* infection contributed 49.4% and others elements host contributed 19.4% of the differences, respectively. Combining these two factors, we assume that *Elaphomyces* in soil are benefit for *Russula* symbiosis.

### Bacterial diversity analysis and analysis of indicator species associated with ECM *Russula* rhizosphere soil

In total, 1494 bacterial operational taxonomic units (OTUs) were distinguished in this study. Compared to type 3, the Chao1 diversity index decreased in types 1 and 2 (Tab. 1). There were significant differences between type 3 and the other two types. However, there were no significant differences between types 1 and 2. The Shannon diversity index showed the same tendency as the Chao1 diversity index (Tab. 1). The bacterial community composition and the abundance of the main bacterial (the over 0.05% bacteria in soil genome) were analyzed (Fig. 3A). About 40-60% of the species in the sample cannot be identified by Illumina MiSeq high-throughput sequencing and the remaining bacterial genera in the sporocarp-producing *Russula* root rhizosphere soil were showed for the three types (Fig. 3A). Compared to the fungi of the root and rhizosphere soil, the bacterial genera variety still increased (Fig. 1A, 2A, 3A).

**Fig. 3.**
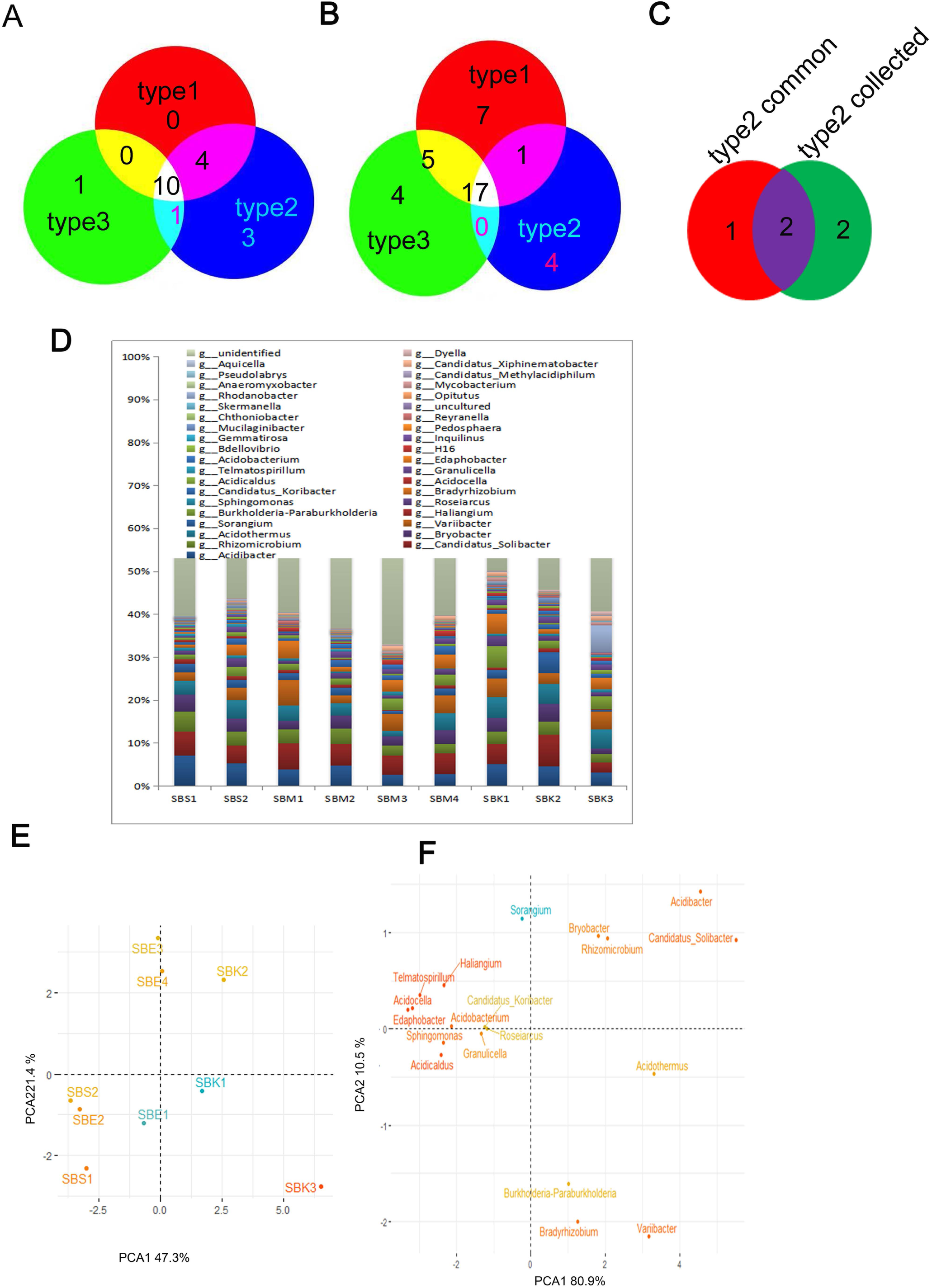
Diversity analysis of bacterial of three types *Russula* Mycorrhizal rhizosphere in natural *Russula* growth areas. A: The top 20 bacterial genus **c**omposition and abundance in type2 *Russula* symbiosis mycorrhizal rhizosphere and the corresponding genus are in the other two types in natural *Russula* growth areas, Fujian, China In the sample name, the first letter s indicates mycorrhizal root soil; first second b stands for bacteria; the third letter different treatment :S indicates sporocarp, E indicates ECM ectomycorrhizal, K indicates control respectively; the fourth number indicates the different biological repetitions of different treatment. B: The Venny mapping of the common genus of the top 20 bacterial genus in three types *Russula* symbiosis Mycorrhizal rhizosphere C: The Venny mapping of the collection genus of the top 20 bacterial genus in three types *Russula* symbiosis Mycorrhizal rhizosphere D: The Venny mapping of the common genus and the collection genus of the top 20 bacterial genus in type2 *Russula* symbiosis Mycorrhizal rhizosphere E: Analysis of 20 bacterial genus role difference of three types Russula symbiosis mycorrhizal rhizosphere on top 20 bacterial genus of ECM Russula by PCoA F : **A**nalysis of **c**omposition difference of three types *Russula* symbiosis mycorrhizal rhizosphere on top 20 bacterial genus of ECM *Russula* by PCoA In E and F:The scales of the horizontal and vertical axes are relative distances and have no practical significance. The contribution rate is the degree of interpretation, and the hypothetical factors can be evaluated and verified.

The community composition of the top 20 f bacteria in the three *Russula* rhizosphere soil types was analyzed (Tab. S3).We first obtained the common species of the three types after intersecting each type with the Venny mapping tool: 18 species in type 2, 14 in type 1 and 12 in type 3 (Fig. 3B). Then, we intersected the common types of the three types with the Venny mapping tool. At the genus level, the three types shared 10 common genera: *Acidibacter, Candidatus, Rhizomicrobium, Acidothermus, Variibacter, Burkholderia, Roseiarcus, Bradyrhizobium, Candidatus* and *Granulicella*. (Fig. 3B). This showed that 10 genera are common bacteria in Fagaceae-dominant rhizosphere soil. Types 1 and 2 shared 4 common genera: *Bryobacter, Sorangium, Acidicaldus*, and *Acidobacterium*,, in addition to the abovementioned 10 common genera(Fig. 3B). Second, we obtained the collection of the various types: 20 species in type 2, 27 in type 1 and 26 in type 3 (Fig. 3C,Tab S3). Then, the unique genera were analyzed of the unique collection (4 genera) and union (3 genera) in type 2 by the Venny mapping tool, and 2 genera ***Acidocella*** and ***Edaphobacter*** were found (Fig. 3D). In all, *Acidocella, Edaphobacter* may be considered indicator species for sporocarp-producing *Russula*, and *Bryobacter, Sorangium, Acidicaldus*, and *Acidobacterium* can be considered indicator species for *Russula* symbiosis in the bacterial community (Tab. 2). By PCoA based on the top 20 fungal genera, we found that type 2 belonged to quadrant III, left of the vertical axis. Except for one sample, other samples of type 1 belonged to quadrants II or III, left of the vertical axis. Type 3 belonged to the right of the vertical axis (Fig. 3F). We assume that the *Russula* infection and host contributed 47.3% and 21.4%, respectively(Fig. 3F). Analyze the different effects of bacteria species by PCoA, we found ***Acidocella***, *Sorangium, Haliangium, Telmatospirillum*, ***Edaphobacter***, *Acidobacterium, Sphingomonas, Candidatus*_*Koribacter, Roseiarcus, Granulicella* and *Acidicaldus* belonged to quadrant II, III, while the others belonged to quadrant I, IV (Fig. 3E). It shows 11 gengus in quadrant II, III can be divided into one categories by role. All indicator species for sporocarp-producing *Russula* or *Russula* symbiosis in the same categories but *Bryobacter.* We assume that the *Russula* infection contributed 80.9% and others elements contributed 10.5% respectively. Combined with indicator species analysis, we assume that 11 bacterial gengus beneficial to the symbiosis of *Russula*.

### Interaction of fungi and bacteria with ECM *Russula* root and mycorrhizal rhizosphere

By network net, we analyzed the interaction of the top 20 fungal OTUs of *Russula* rhizosphere soil and ECM *Russula* root, including 7 Russula OTUs(MSFH OTU_1, MSFH OTU_3, MSFH OTU_4, MSFH OTU_16, MSFH OTU_21, MSFH OTU_24 and SMFH OTU_1639) (Table S7), five indicator species(MSFH OTU_19 *Elaphomyces*, MSFH OTU_9 *Tomentella*_sp, MSFH OTU_655 *Tomentella*_sp, MSFH OTU_6 *Elaphomyces*, and MSFH OTU_5 *Tomentella*_sp).

*Russula*, which interacted with other species in positive ways, was the representative and contributed to the main ECM in the community. For example, the interaction result predicted that many fungi had a positive correlation with *Russula* (Fig.4A, Table S4), MSFH OTU_19 (*Elaphomyces*) and MSFH OTU_5 (*Tomentella*_sp) with the *Russula rosea*; MSFH OTU_21, MSFH OTU_655 (*Tomentella*_sp) and MSFH OTU_5 with Russula sp. In all, *Elaphomyces*_sp, *Tomentella*_sp have a positive correlation with *Russula*, combining the previous results that *Elaphomyces* and *Tomentella* were considered indicator species for *Russula* symbiosis in the *Russula* rhizosphere based on the top 20 genera (Tab. 2). Therefore, we further assume that the indicators *Elaphomyces* and *Tomentella* have a positive correlation with *Russula* symbiosis.that Meanwhile, we analyzed the relationship of the top 20 bacterial OTUs in *Russula* rhizosphere soil by Network net. Only 15 bacterial OTUs were included in the network net (Fig. 4B, Table S5). Among the top 15 bacterial OTUs, there is 1 OTUs SBH OTU_70 *Sorangium* belonging to indicator species. The network net suggested that SBH OTU_70 was positive with SBH OTU_16 (Acidobacteriaceae), 24(Caulobacteraceae), 27(Solibacteraceae), 69(Solibacteraceae). Another indicator species *Acidobacterium* was’t within network net. Therefore, we further assumed that *Sorangium* and other OTUs (Acidobacteriaceae, Caulobacteraceae, Solibacteraceae) have a positive correlation with *Russula*.

**Fig. 4.**
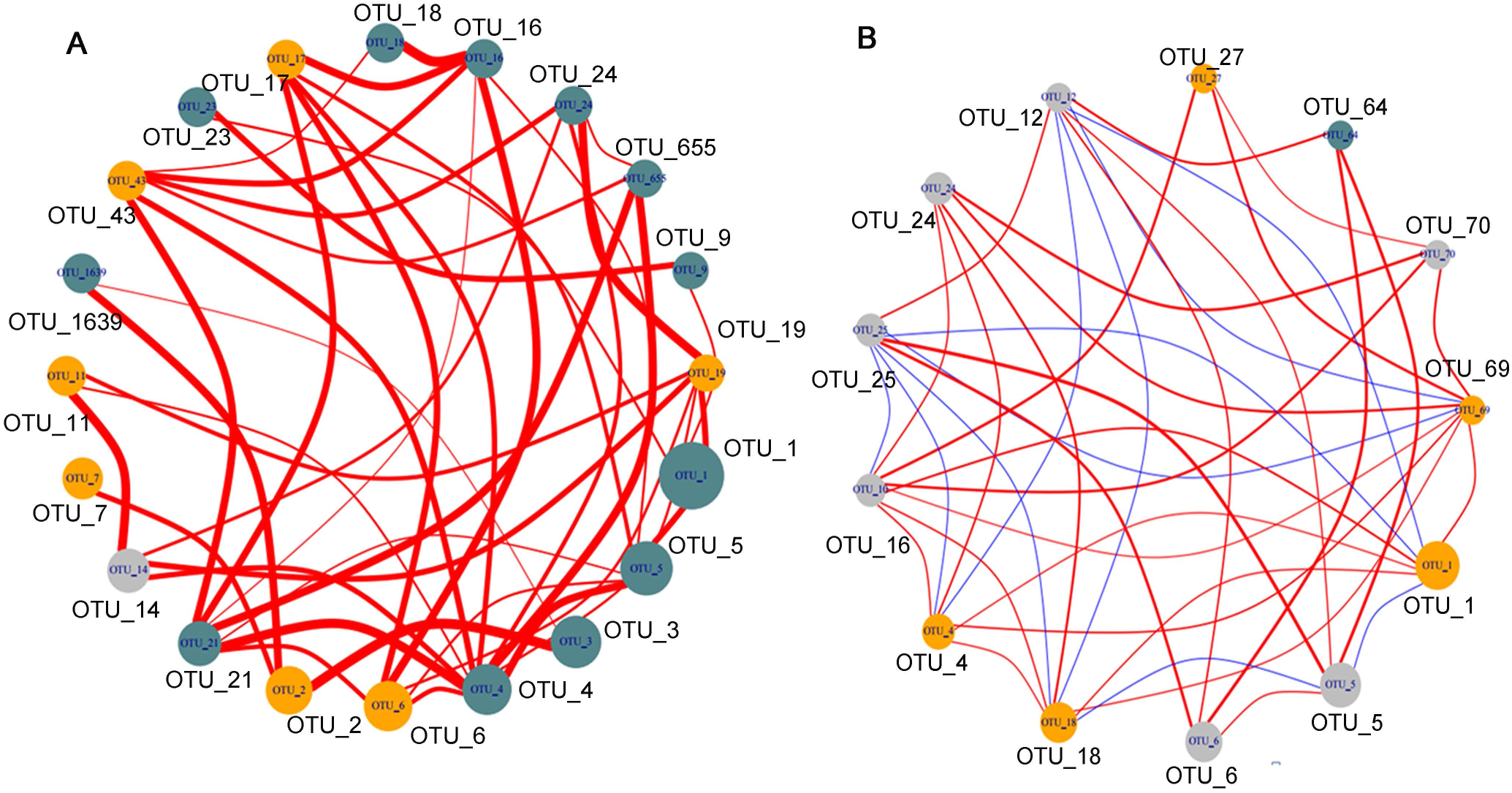
Interaction of indicator species with the top 20 fungi and bacterial OTUs in community of *Russula* mycorrhizal. A: Interaction of fungi in *Russula* ECM root and *Russula* mycorrhizal rhizosphere B: Interaction of bacterial in Russula mycorrhizal rhizosphere In A and B: The size of the dots represents the abundance, the color of the dots represents the phylum (In A : Orange represents p__Ascomycota, blue represents p__Basidiomycota, gray represents p__Mortierellomycota; in B: gray represents p__Proteobacteria, Orange represents p__Acidobacteria, blue represents p__Actinobacteria), the thickness of the lines represents the magnitude of the correlation, and the red line indicates a positive correlation, the blue color indicates a negative correlation.

## DISCUSSION

The complete exclusion of other species by any single species could be prevented by either fluctuation in the environment, by seasonal root production or by the presence of microbiome competitive networks, the situation in which no single species is competitively superior to all other species. So, we usually obtain 1 ECM species or 2 ECM species by Sanger sequencing of the root sample with basidiomycete-specific primers. The introduction of high-throughput sequencing techniques, metagenomics or environmental genomics has provided new information on ECM fungal communities by ‘barcodes’ of ITS regions in several biomes/ecosystems, e.g., truffle grounds (57) and ECM roots in the Svalbard(58). In this study, we found that *Russula* cannot successfully infect roots if the amount of *Russula* in the mycorrhizal rhizosphere is less than 10% by MiSeq sequencing (Table 1, part data not shown). Although ECM fungi infection have been detected in nonproductive plots, the amount was not sufficient to shift from vegetative growth to fruit body. *Russula* can form a fruiting body, only when *Russula* infection could be detected by Sanger sequencing of ECM roots, contemporary the relative abundance of its DNA was greater than 60% by MiSeq sequencing. Compared to control, we found significantly more *Russula* DNA in *Russula* sporocarps rhizosphere soil or *Russula* symbiosis rhizosphere soil, above 15% amount of the metagenomics of rhizosphere soil. Zampieri et al.(2012) (49) detected significantly more *T. melanosporum* DNA in truffle productive plots of soil. Zhou et al. (2001) (50) also demonstrated that the development of *S.grevillei* sporocarps is correlated with that of extraradical mycelia, which are distributed in a narrow area. So, we suggest that to the type 2(sporocarps-producing), there is congruence of the above-sporocarps and belowground root or mycorrhizospheres soil DNA for *Russula*. However, interestingly, in type 3(uninfected), the amount of *Russula* in the soil is higher than in the root. This may indicate that in the natural growth area of *Russula*, there are a large number of *Russula* propagules in the soil. Whether the *Russula* propagules naturally in the soil of the natural growth area can colonize to the host or not depends on other factors.

Fewer species were detected in *Russula* productive root than in nonproductive root, which is consistent with French *truffle* grounds. *T. melanosporum* grounds also have shown a reduced fungal biodiversity, a reduced presence of both ECM Basidiomycota (57). Truffle-colonization reduced the abundance of some fungal genera surrounding the host tree, such as Acremonium (55). *Russula* fruiting body decreased the Chao richness bacterial index and Shannon diversity of bacterial index in rhizosphere soil. Simultaneously, *Russula* fruiting body decreased the Chao richness fungi index but no effect on Shannon diversity of fungi index in rhizosphere soil. Warmin (2009)(40) found bacteria adapted to the mycospheres of three or more or just one ECM fungal species was defined as specific selective bacterial. So, we suggest the diversity decreased phenomenon may be related with the *Russula* sporocarp selection effect, especially to rhizosphere soil bacterial. Temperate forests are generally dominated by Fagaceae, and species in these plant families form mycorrhizae with various phylogenetic clades of ectomycorrhizal fungi(8, 11, 59). Members of the *helotialean* group have recently been identified as the dominant species in the roots of Fagaceae trees in the temperate and subtropical forests of Japan (60, 61). In our study, we found that 3 ECM, *Russula, Tomentella*, and *Lactarius*, are the main members in the natural Fagaceae *(Quercus glauca* and *Castanopsis hainanensis*)-dominant *Russula* ground in all the types. This result is consistent with the preference of *Russula* and *Lactarius* for the Fagaceae host. In *Castanopsis*-dominant forest in Japan (62), there are ECM fungi such as *Amanita* (Amanitaceae), *Boletus* (Boletaceae), *Tylopilus* (Boletaceae), *Cortinarius* (Cortinariaceae), *Inocybe* (Inocybaceae), *Lactarius* (Russulaceae), and *Russula* (Russulaceae). The dominant ECM lineages of *Quercus liaotungensis* were *Tomentella, Thelephora, Cenococcum, Russula, Lactarius* and *Inocybe*(63). The ectomycorrhizal *Russula, Lactarius, Cortinarius, Tomentella, Amanita, Boletus* and *Cenococcum* were dominant in the Quercus serrate plot (11). *R. vinosa* grows in tropical and subtropical evergreen forests in southern China dominated by trees of *Castanopsis* spp., and *R. griseocarnosa* grows in forests with Fagaceae (13). Given commercially harvested truffles can establish ectomycorrhizal relationships with different woody host species, and many different combinations of truffle and host species mycorrhizal seedlings are produced in nurseries (64). Our study shows that *Russula* inoculation may establish satisfactory ectomycorrhizal relationships with two indigenous tree species, *Quercus glauca* and *Castanopsis hainanensis.*

Healthy ECM can support a wide variety of organisms, including a diverse array of fungi other than the dominant ectomycorrhizal symbiont. Some pathogenic fungi, including *Ilyonectria* and *Podospora*, and other competitive mycorrhizal fungi, such as *Hymenochaete*, had significantly lower abundance in the *T. borchii*-inoculated root and *Trechispora* and *Humicola*, which were more abundant in the *T. borchii*-inoculated root. There are some notable examples of associations among *suilloid* fungi and members of the Gomphidiaceae occur within ectomycorrhizal roots (27). Olsson considered *Gomphidius roseus* acted as a parasite of *Suillus bovinus*, the former never occurs without the latter(27), Based on MiSeq sequencing analysis of the top 20 ectomycorrhizal fungi in *Russula* root, we found *Acremonium*; *Cladophialophora* are associated with ECM *Russula*-Fagaceae roots, and *Boletus* was in association with sporocarp-producing *Russula*. Meanwhile, based on the top 20 genera in *Russula* rhizosphere soil, we found that *Elaphomyces*, and *Tomentella* in association with *Russula* symbiosis; *Dacryobolus* were associated with sporocarp-producing *Russula*. The result will help to find and develop PGPF for *Russula* symbiosis or sporocarp-producing.

Many studies have addressed the role of soil bacteria in establishing the symbiotic relationship between plants and mycorrhizal fungi (43, 65). We found differences in fungal genus among the three ECM Russula types in rhizosphere soil (Table S 3, Table 2).The interactions of bacteria with the dense hyphal network underneath fungal fruiting bodies have also been addressed (44, 66). Differences in bacterial communities associated with the mycorrhizospheres of *Suillus bovinus*- and *Paxillus involutus*-colonized plants were detected early (67, 68). Frey et al.(1997)(69) reported that specific *Pseudomonas fluorescens* prefer *Laccaria laccata. P.fluorescens* and *Burkholderia terrae* are exclusively found in the mycosphere soil of *Laccaria proxima* (30, 40). Boersma(2009)(36) found that the mycospheres of basidiomycetous fungi indeed exerted a universal selective effect on the *Pseudomonas* community (i.e., *Pseudomonas poae, P. lini, P. umsongensis, P. corrugata, P. antarctica* and *Rahnella aquatilis*); as well as species-specific selective (i.e. *P. viridiflava* and candidatus *Xiphinematobacter americani*). For the selection of the bacteria family *Sphingomonadaceae* by the mycorrhizal fungi *L. proxima* and *R. exalbicans*, the mycosphere effect was most prominent in the latter (36). The mycosphere-isolated bacterium *Burkholderia terrae* has been shown to protect its fungal host *Lyophyllum* sp. from several antifungal agents, such as metabolites produced by *P. fluorescens*, as well as from the antifungal agent cycloheximide (70). Some studies have concentrated on mycorrhization helper bacteria (MHB) in facilitating mycorrhizal colonization of roots in forest nursery environments(43, 71). Rich in bacteria in the mycorrhizal roots, mycorrhizosphere soil and peridium of desert truffles may be used to increase the survival and mycorrhization in the desert truffle plant production system at a semi-industrial scale(72). The associated bacteria of *Truffle brûlés* are *Pseudomonas* and *Flavobacteriaceae* (73). To achieve successful reforestation, PGPR and MHB were screened to improve the establishment of *Lactarius deliciosus*-Pinus sp. symbiosis(74). In our study, *Acidocella, Bryobacter, Sorangium, Acidicaldus, Edaphobacter* and *Acidobacterium* were indicator species for *Russula* symbiosis in the bacterial community. we also found ECM universal selective *Sphingomonadaceae* work with *Russula* species-specific selective *Acidocella, Sorangium, Acidicaldus, Edaphobacter* and *Acidobacterium* in the soil bacterial community of *Russula* symbiosis (Fig. 3E).Further, by network analysis, *Acidobacteriaceae, Sorangium* and *Acidobacteria* had a positive correlation with *Russula*. There may be further instruction to provide a wide collection of these bacterial associates of *Russula* and to develop *Russula* MHB or Fagaceae PGPR.

Only a few edible ectomycorrhizal fungal species, such as black truffles or saffron milk caps, can be produced in manufactured orchards. To date, *Russula* fruit bodies are uncultivable. Five *Russula* were used to inoculate *Shorea parvifolia* seeding successfully (75), and one *Russula* was used to inoculate *Quercus garryana* seeding successfully(76). Our results will provide instruction to specifically isolate the fungi or beneficial rhizosphere microbes associated with ECM *Russula* from the root or mycorrhizal rhizosphere. Next, we will further need to isolate and culture the microbial communities of the *Russula* root or mycorrhizospheres soil in three types, which will supplement our research findings. The inoculation of these microbes can stimulate establishment and will help to enhance plant growth and promote a change in infected frequency.

### Conclusion

The amount of *Russula* DNA is positively correlated with fruiting body and *Russula* mycorrhizae based on the metagenomics of *Russula* root and soil. Fewer fungi species were detected in *Russula-* infected and *Russula* sporocarp root. Fewer bacteria and fungi species were detected in *Russula-* infected and in *Russula* sporocarp rhizosphere soil. *Boletu*s is considered as indicator species in Russula-Fagaceae root (*Quercus glauca* and *Castanopsis hainanensis*) for sporocarp-producing *Russula*. The *Russula* sporocarp rhizosphere fungi *Dacryobolu* and *Russula* sporocarp rhizosphere bacteria *Acidocella* is considered as indicator species for sporocarp-producing *Russula*. On the other hand, a number of taxa within *Acremonium* and *Cladophialophora* were identified in Fagaceae root in *Russula* symbiosis. The *Russula* mycorrhizal rhizosphere fungi *Tomentella*, and *Elaphomyces* and the *Russula* mycorrhizal rhizosphere bacteria *Acidocella, Bryobacter, Sorangium* and *Acidobacterium* occurred more frequently in association with the ECM genus *Russula*. Understanding the ectomycorrhizal fungal communities inhabiting natural *Russula* growth areas may give us clues about the dynamics of the targeted *Russula* and the possibility of identifying mycorrhizal fungal species that are good indicators of successful *Russula* semicultivation. This reseach may provide novel targeted strategies to improve the establishment of *Russula*-Fagaceae sp. symbiosis and improve *Russula* ascospore productivity and sustainability.

## MATERIALS AND METHODS

### Sampling

Our study was conducted in areas of *Russula* growth in Jianou, Fujian province, China, in which species of the genus *Russula* are well represented. The dominant trees of the areas are *Quercus glauca* and *Castanopsis hainanensis*. The herb layer is composed of *Podophyllum peltatum, Panax stipuleanatus* and *Saxifraga stolonifera*. Fine roots were excavated 2-3 m from the trunk of an adult *Russula* symbiotic tree(Fig. S1A). In addition, another type of fine roots that was clearly connected to *Russula* sporocarp by extraradical mycelia was collected(Fig. S1B,C). We collected three 15-30 cm fine-root segments (containing 100-200 root tips) at a depth of 20 cm (77). Roots from a single tree were pooled into a single plastic bag. More than 200 cm^3^ of rhizosphere soil was collected around the root samples for analyses. All root and soil samples were stored in a cooler containing several ice bags and transported to our laboratory within 24 h for subsequent analysis.

### DNA extraction, amplification, and sequencing

The collected root samples were washed carefully with tap water. Root tips were preserved in a plastic centrifuge tube at −20°C before DNA extraction. DNA samples were pulverized using liquid nitrogen. Total genomic DNA was extracted from ECM root tips using a modified cetyltrimethylammonium bromide method, which was modified according to Wang et al. (63)(2017). Total genomic DNA of corresponding rhizosphere soil was extracted using a Fast DNA SPIN Kit (MP Bio) for soil according to the kit operation steps.

First, the entire range of fungal ITS sequences was amplified from roots using the ECM Basidiomycetes-specific high-coverage primer polymerase chain reaction (PCR) with the primer pair ITS-1F (CTTGGTCATTTAGACGAAGTAA) and ITS-4B (CAGGAGACTTGTACACGGTCCAG) (M. Gardes, 1993), and traditional Sanger sequencing was performed. The sequences were BLAST against the UNITE database/NCBI database (http://www.ncbi.nlm.nih.gov), and taxonomy was assigned to species or genera using sequence similarity criteria of ≥97% for species and 90–97% for genera. Furthermore, DNA of root tips identified as *Russula* and the corresponding *Russula* symbiosis rhizosphere soil were subjected to Illumina MiSeq high-throughput sequencing of ITS sequences to investigate their associated microbiomes, while DNA of root tips with no mycorrhiza was used as a control. The MiSeq sequences were edited, manually corrected in BioEdit 7.0.8 and clustered into species-level operational taxonomic units (OTUs) at 97% sequence similarity for species delimitation using PlutoF (http://unite.ut.ee) in UNITE(80). If no match was found in the PlutoF system, any resulting OTU assignments were individually checked by BLAST against the UNITE database/NCBI database (http://www.ncbi.nlm.nih.gov).

To identify the fungi composition, we analyzed 18S ITS1-ITS4 DNA sequences of ECM rhizosphere root and soil samples using phylogenetic methods, operational taxonomic unit (OTU) delimitations and ordinations to compare species composition in various types of ECM *Russula*. To identify the bacterial composition, we analyzed 16S V3-V4 DNA sequences of ECM rhizosphere soil samples using phylogenetic methods, operational taxonomic unit (OTU) delimitations and ordinations to compare species composition in various types of ECM *Russula*.

### Statistical analyses

Chao1 (species richness index) and Shannon (microbial diversity index) indices were analyzed by 97% OTU similarity. ECM fungal richness and the microbial diversity index of each Russula root tip and the corresponding rhizosphere soil of *Russula* symbiosis were calculated using the vegan package and compared by one-way ANOVA. PCoA analyses were based on any distance other than the Euclidean distance using abundance and presence-absence data of the top 20 genera in three types. Differences in community composition among *Russula*-infected samples were visualized by PCoA (79).

### Micro-Community diversity analysis and indicator species analysis

Taxonomic analyses were generated based on the community species abundance (each was above 0.05% of all reads) using Microsoft Excel. The mian fungal genus patterns were determined by taxonomic analysis at different ECM *Russula* mycorrhizal roots. The main fungal and bacterial genus patterns were also observed in the *Russula* mycorrhizal rhizosphere. To assess *Russula* preference, aspects indicator species analysis was carried out by comparing the top 20 genera of three types of root micro-communities. Indicator species of the community of the top 20 genera based on the species presence in the *Russula* sample and absence in the no-*Russula* sample were analyzed by Venny drawing tools (https://bioinfogp.cnb.csic.es/tools/venny/index.html).

### Network analyses

The relationships of *Russula* and other fungi in the *Russula* community were analyzed by Spearman correlation of the absolute top 20 most abundant in all the samples using the igraph, psych software package. Network nets constructed on the interrelationship result with P>0.05 or |R|<0.4 were filtered. The relationships of bacteria in the *Russula* mycorrhizal rhizosphere were also analyzed by Spearman’s correlation.

## SUPPLEMENTAL MATERIAL

Supplemental material for this article may be found at

**SUPPLEMENTAL FILE 1**, TIFF file, 0.48 MB.

**SUPPLEMENTAL FILE 1**, PDF file, 0.22 MB.

**SUPPLEMENTAL FILE 1**, PDF file, 0.22 MB.

**SUPPLEMENTAL FILE 1**, PDF file, 0.22 MB.

**SUPPLEMENTAL FILE 1**, PDF file, 0.17 MB.

**SUPPLEMENTAL FILE 1**, PDF file, 0.13 MB.

## ACKNOWLEDGMENTS

This work was supported by Guiding Program for Science and Technology Department of fujian Province(2017N0005).

---

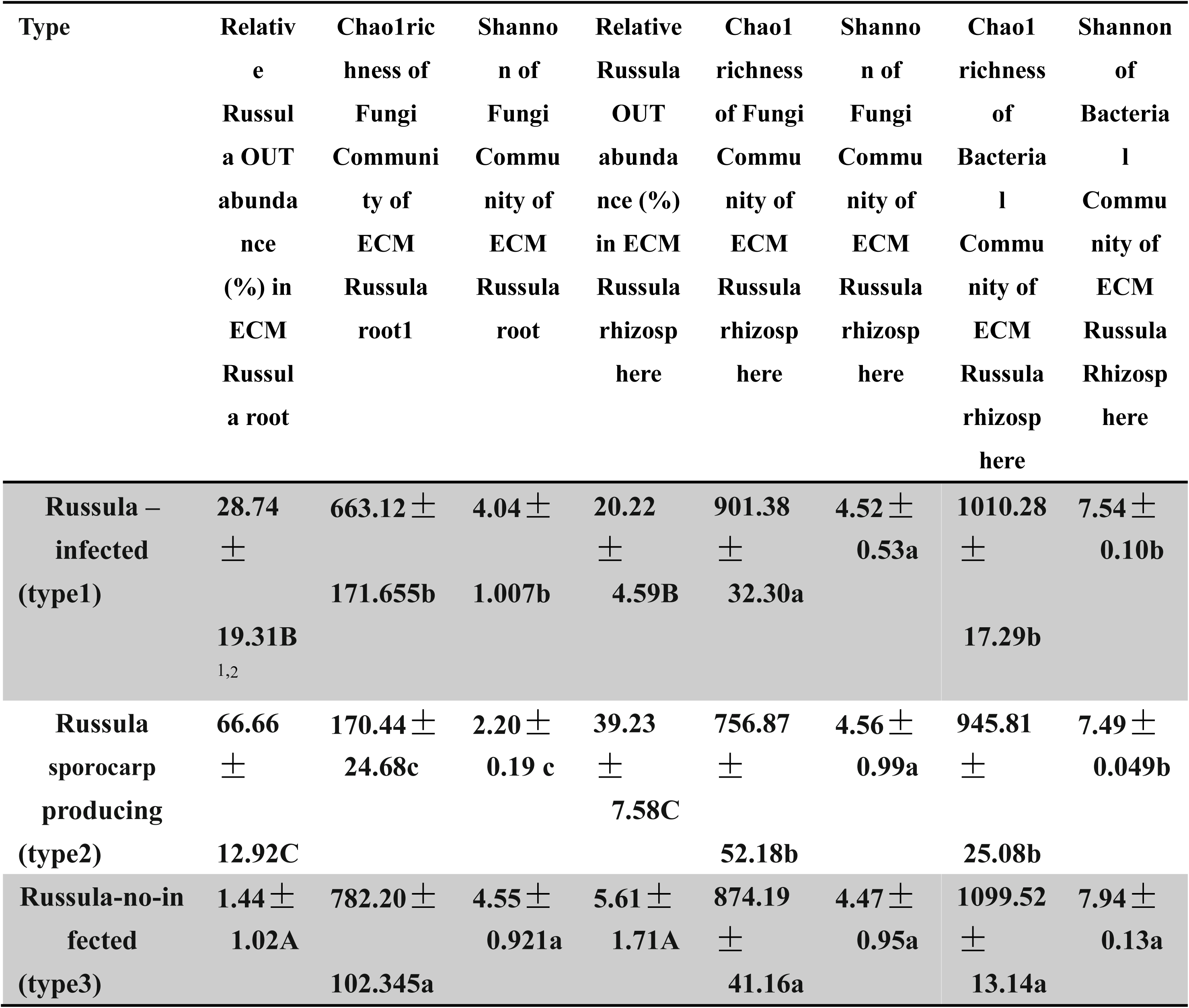
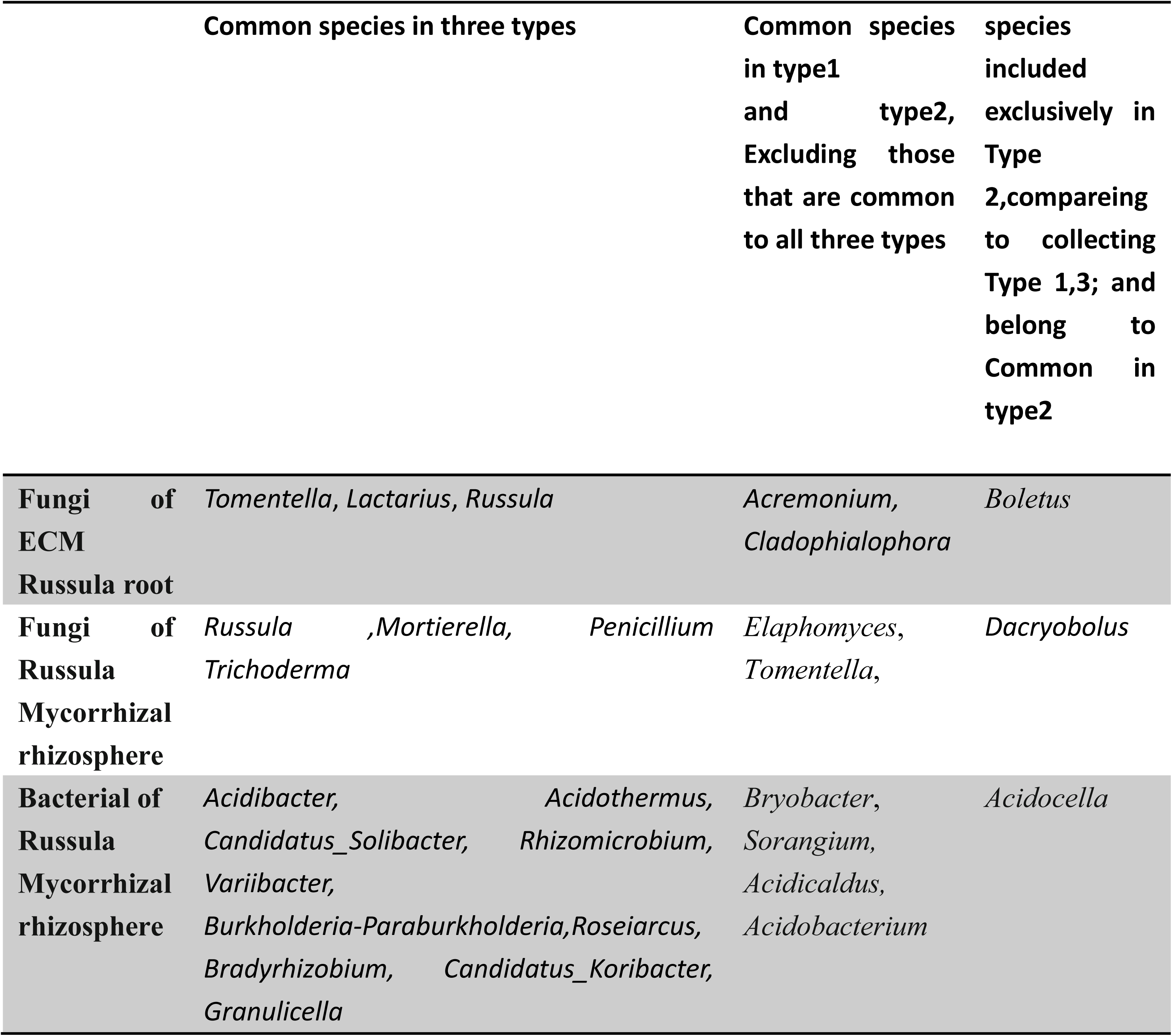

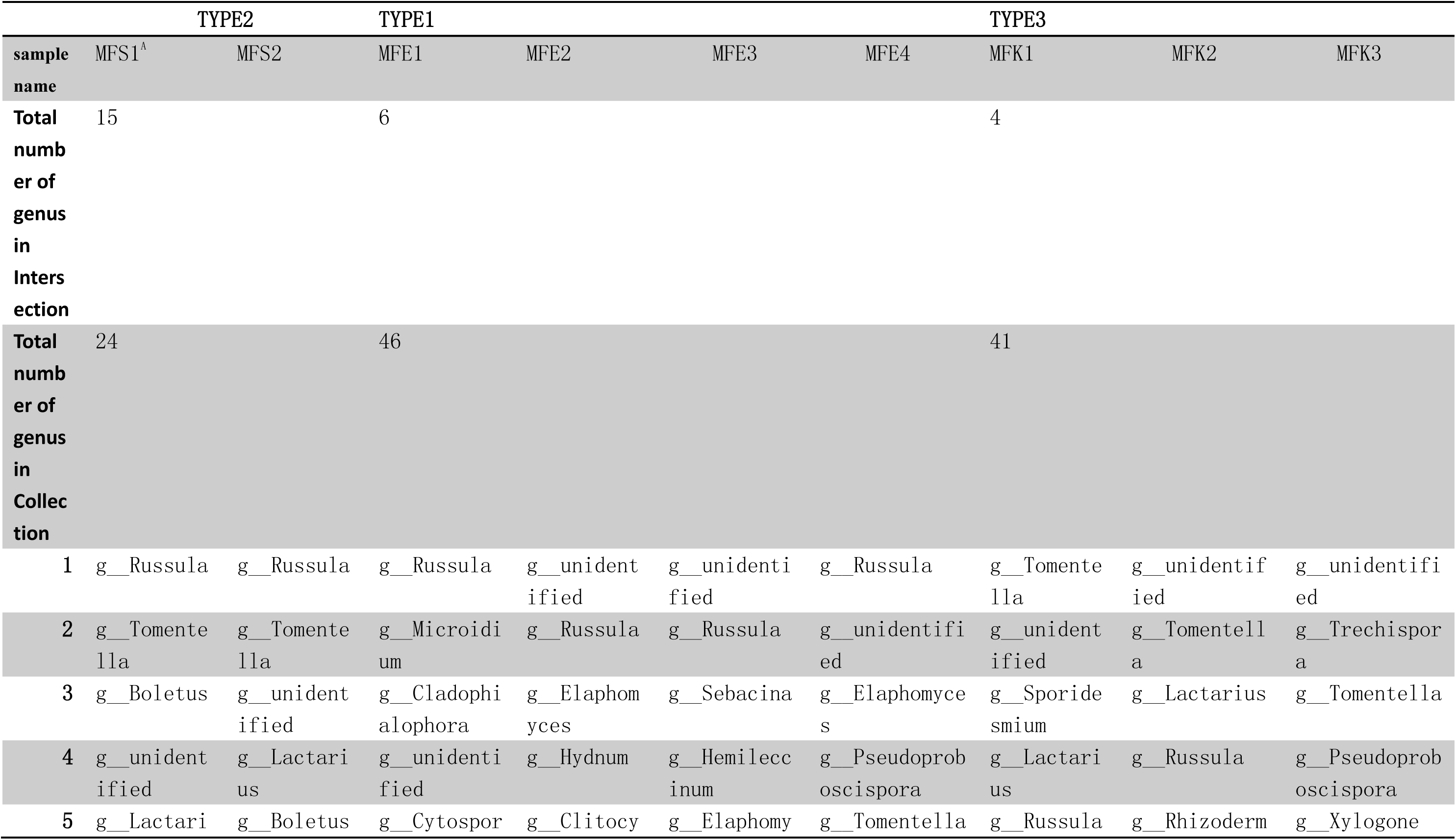
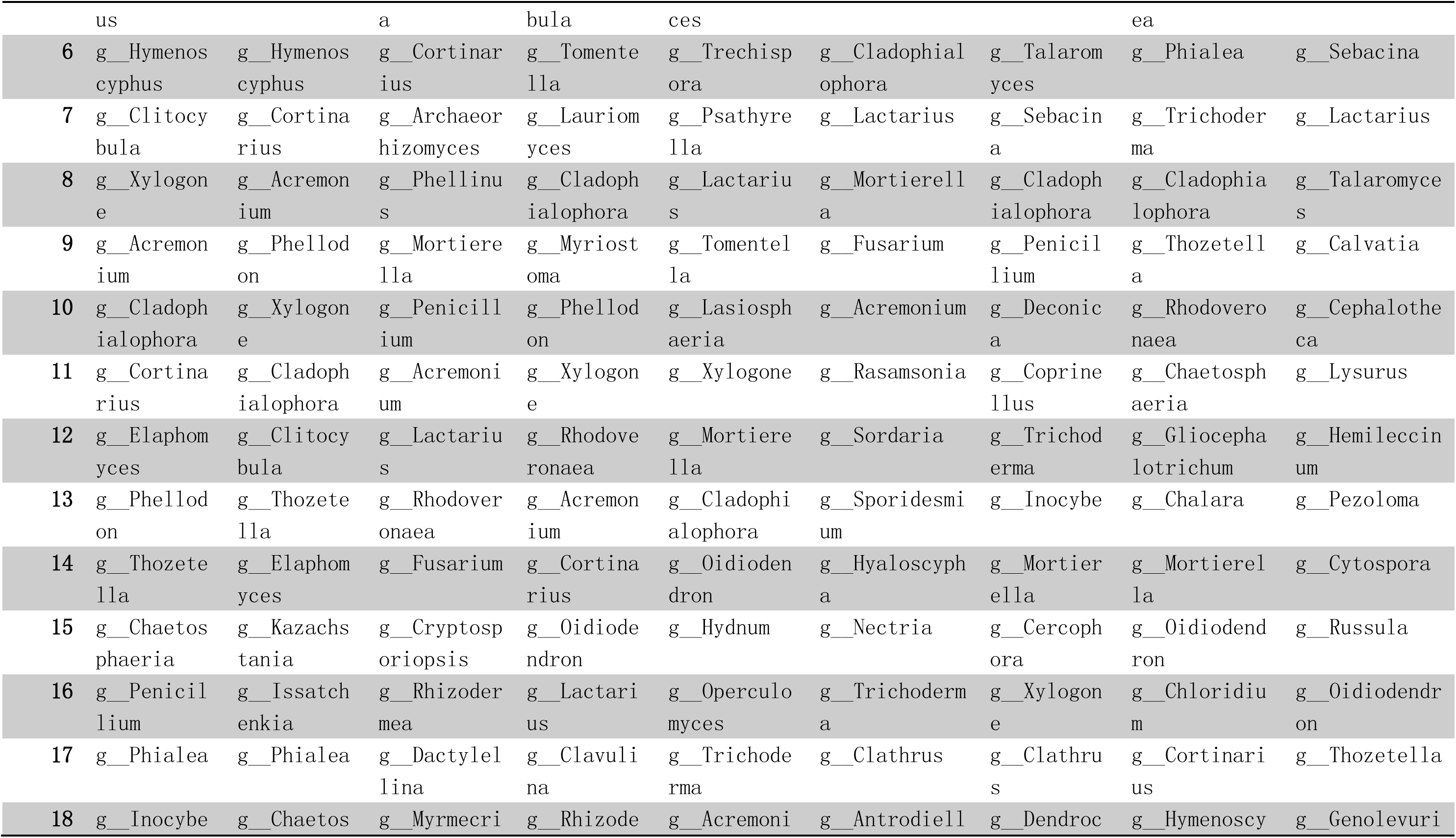
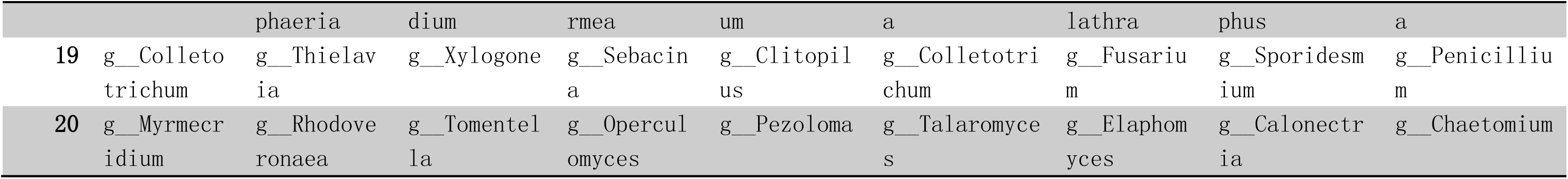

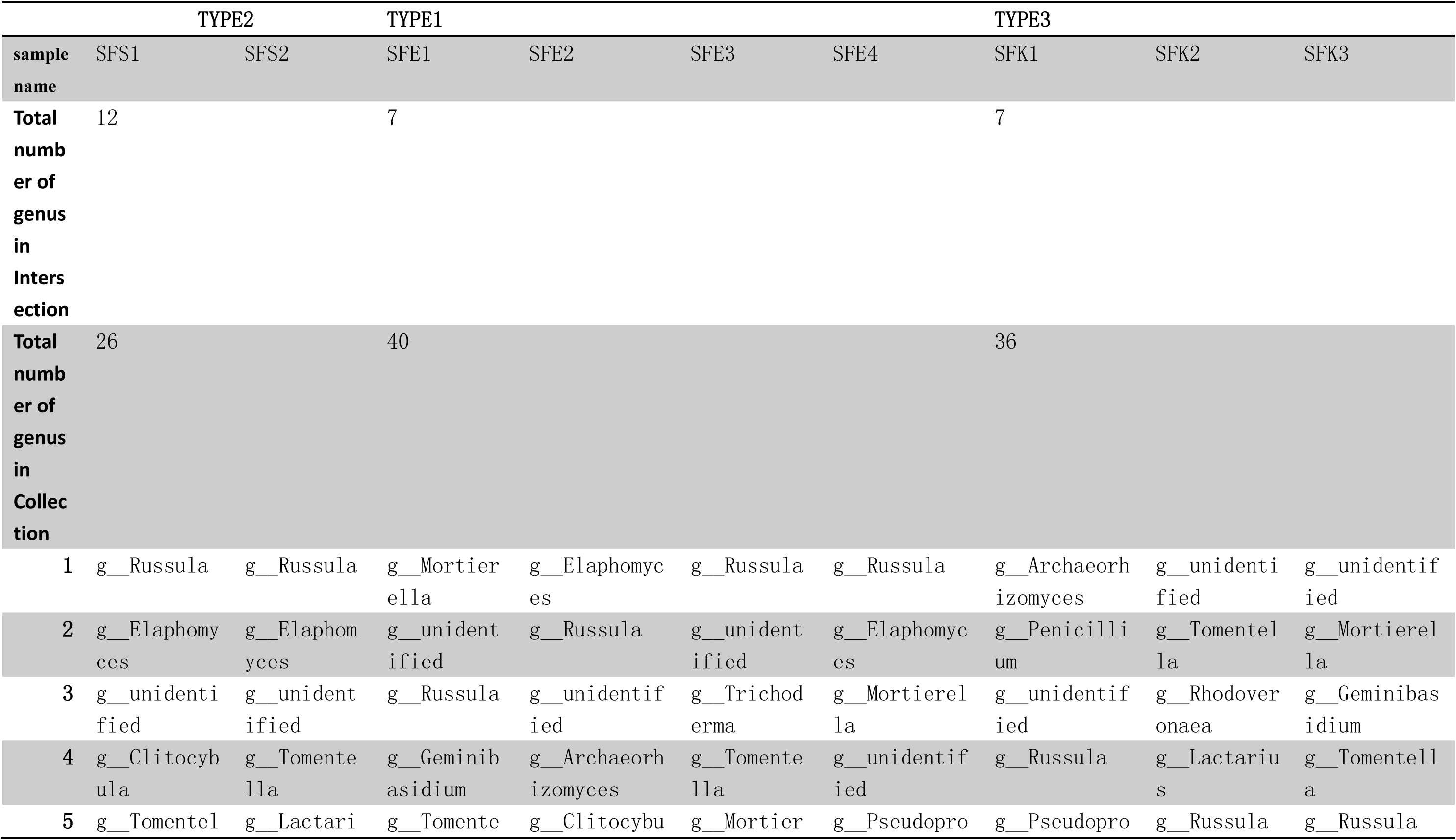
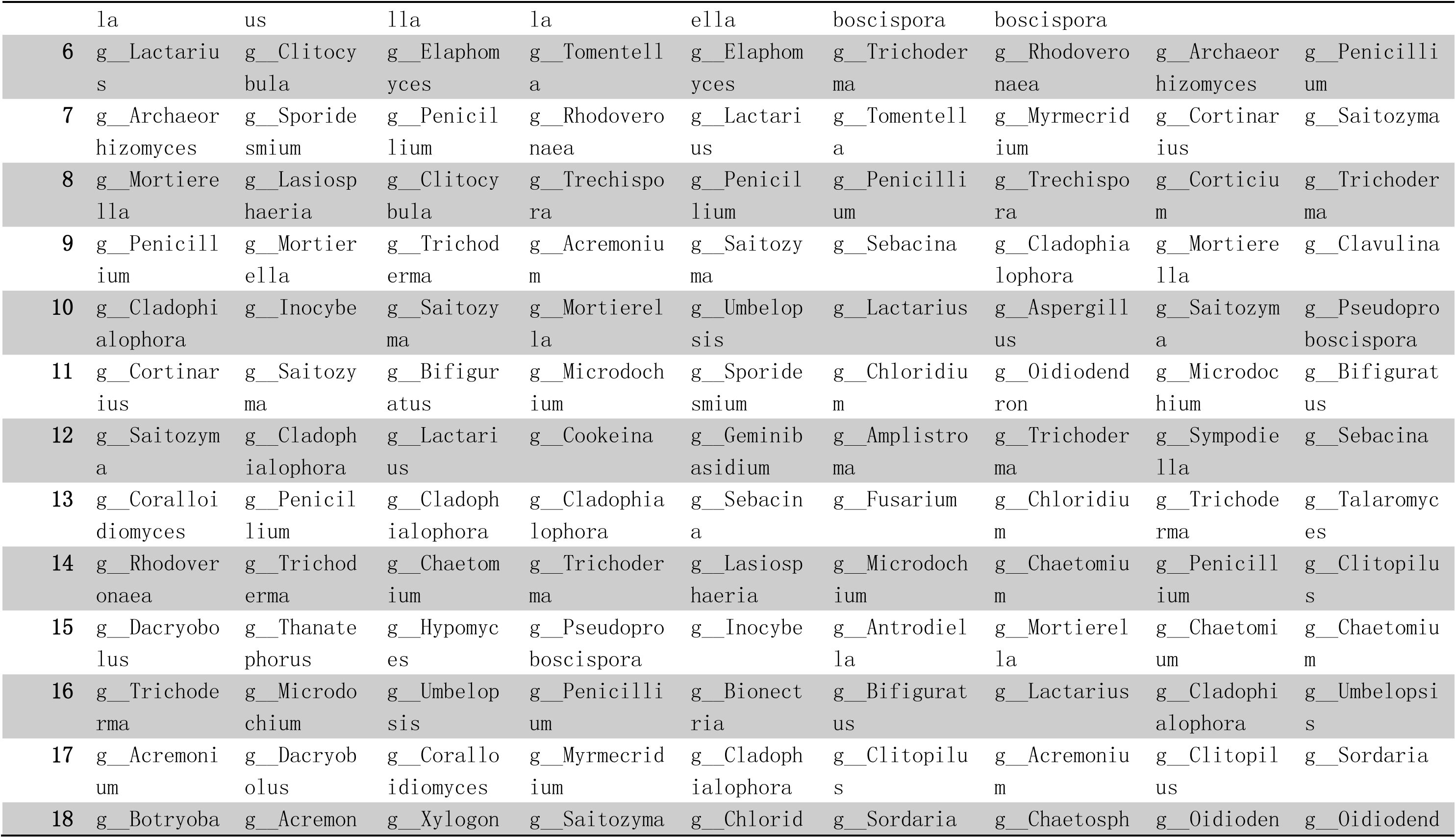
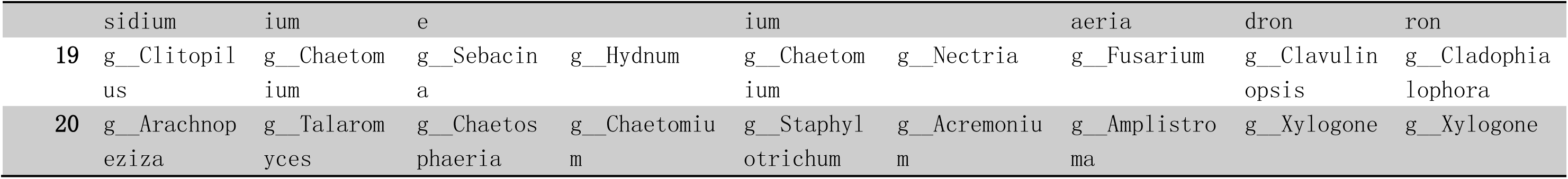

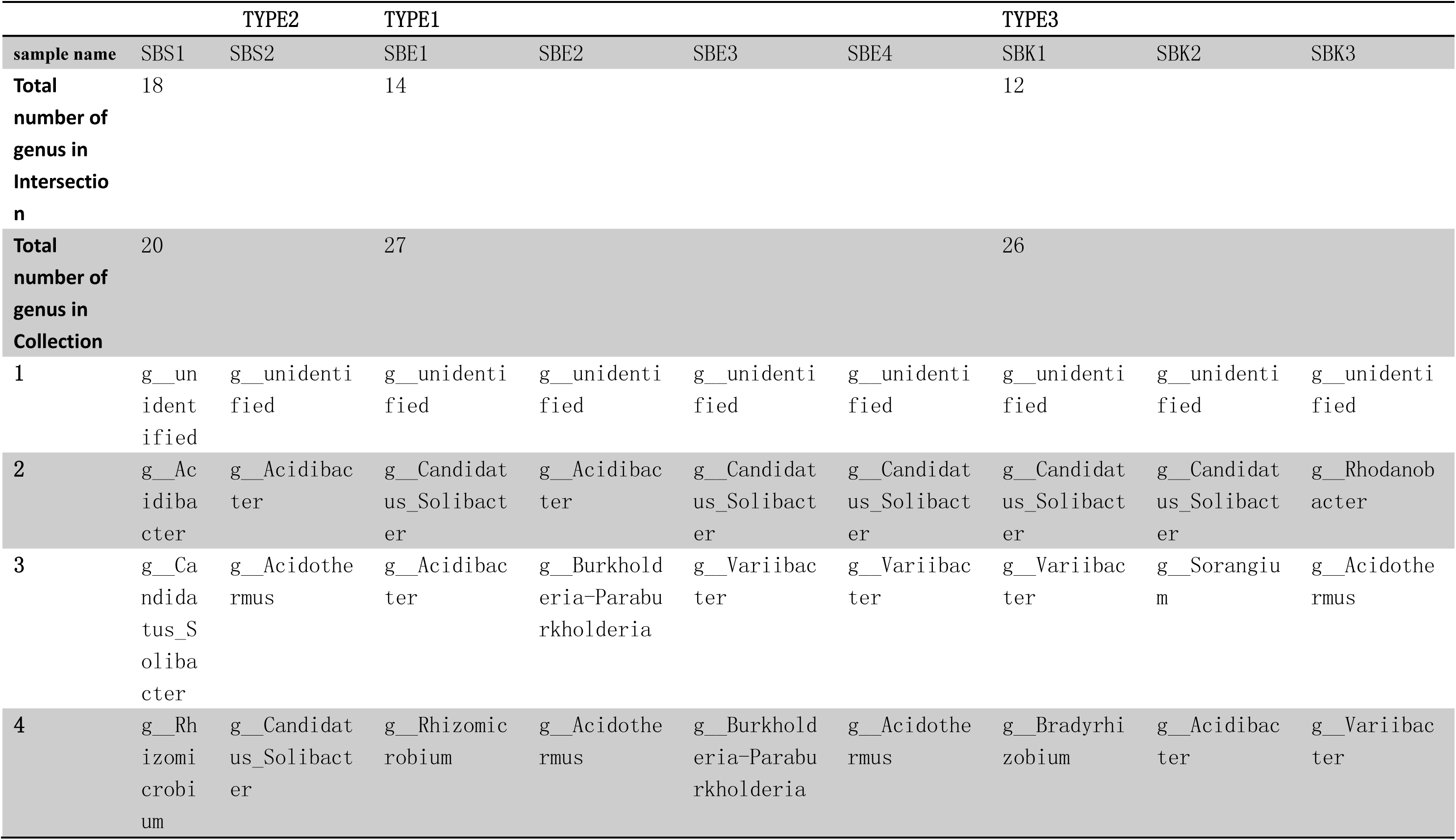
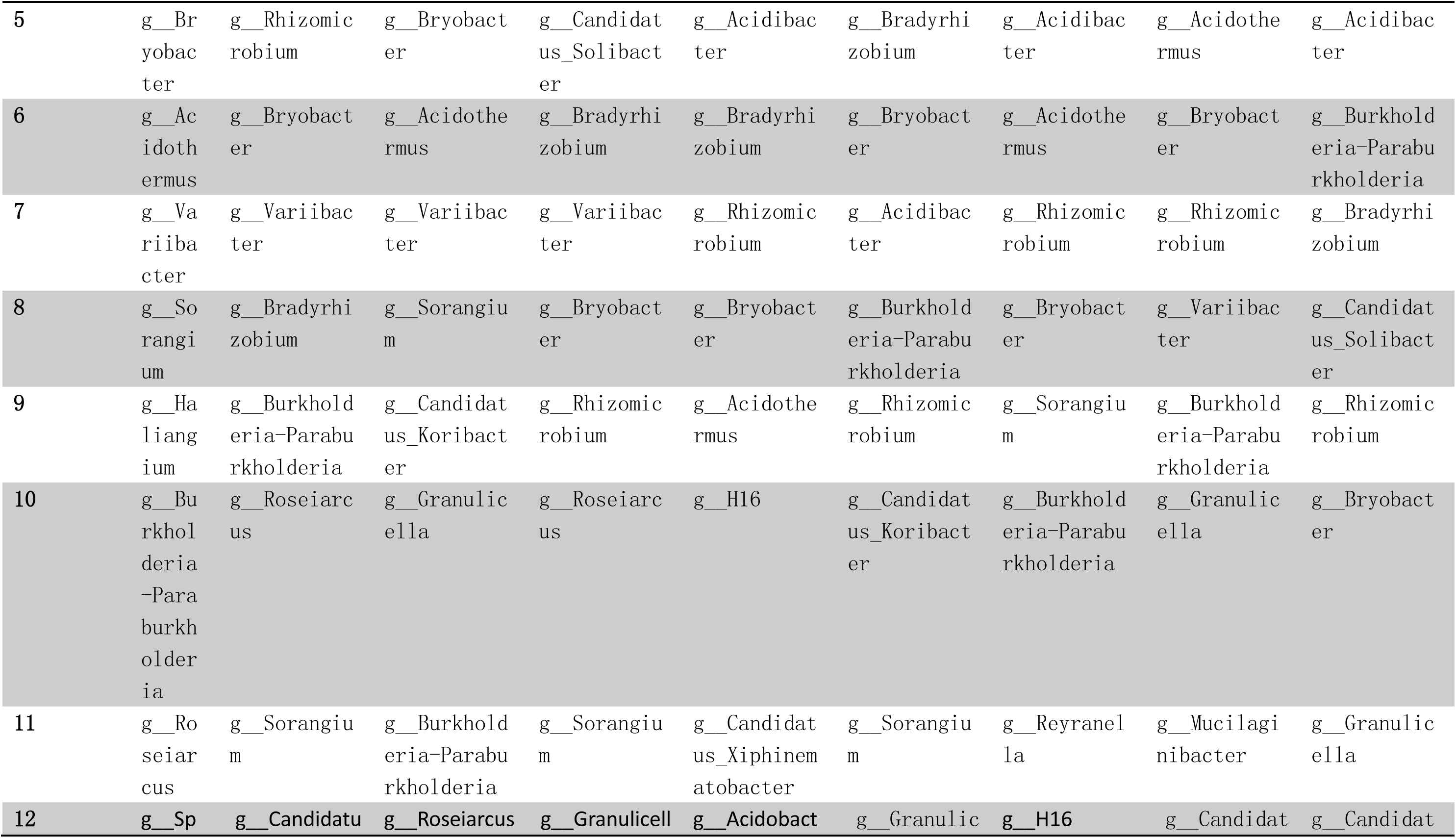
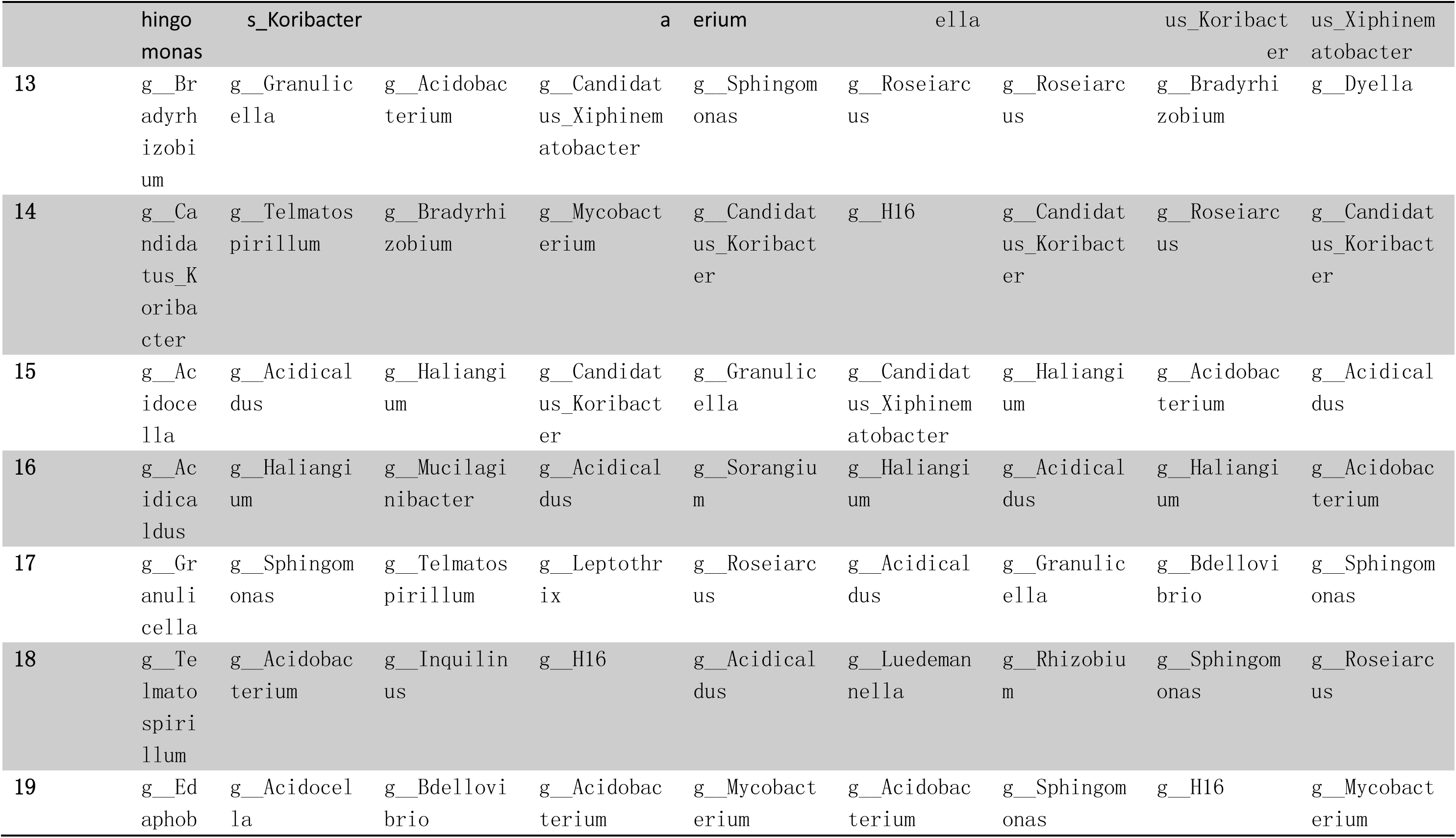
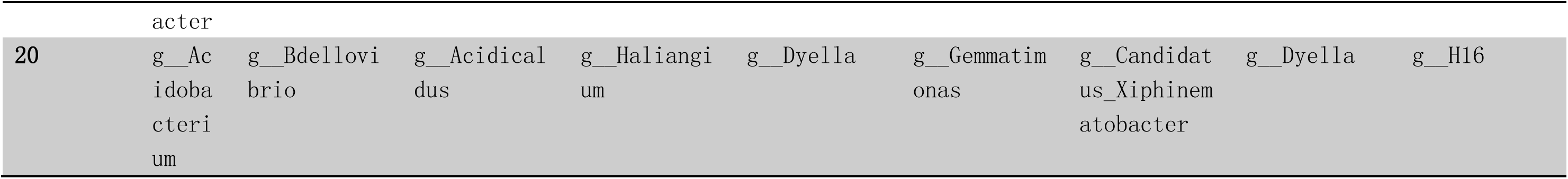

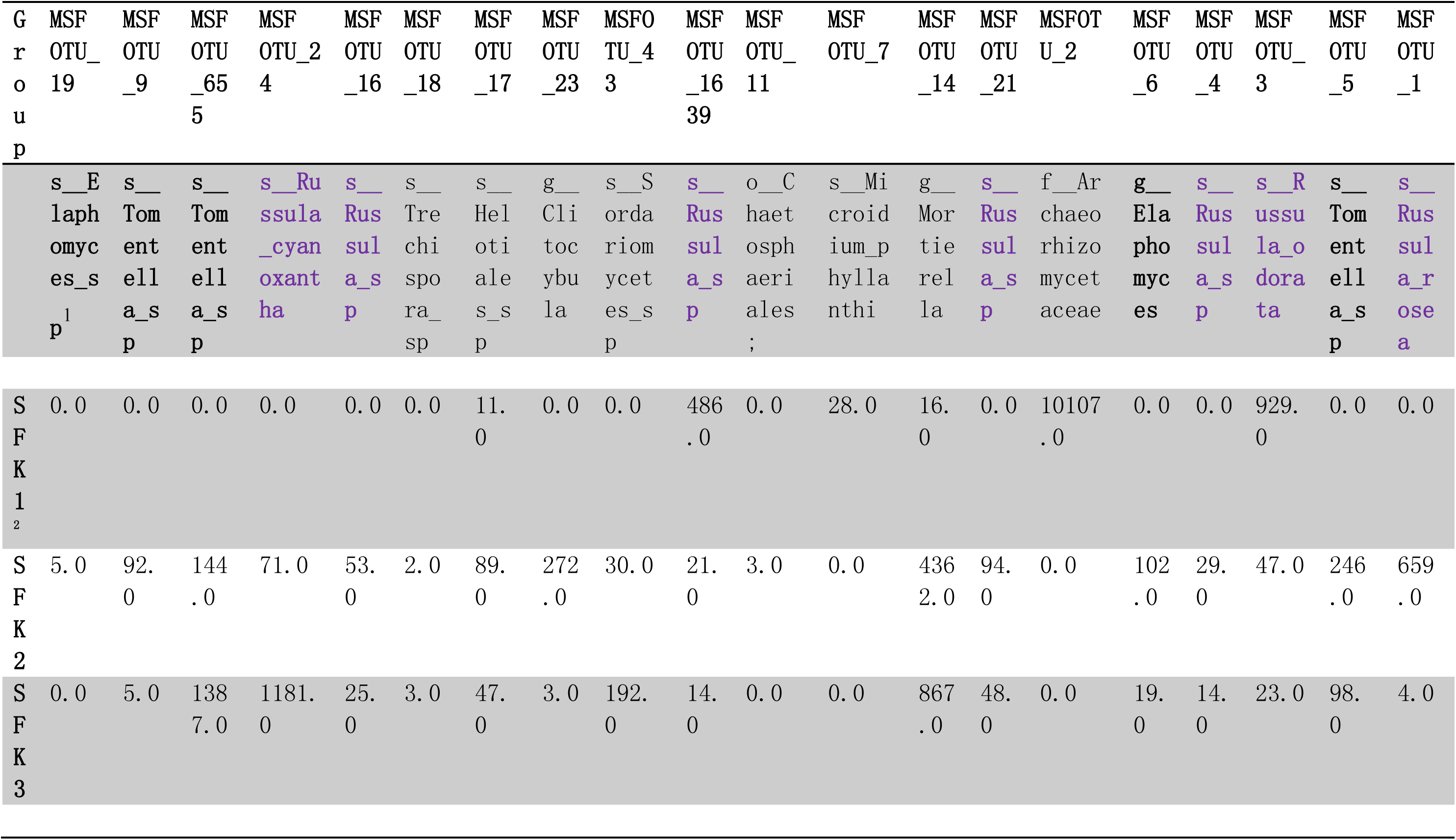
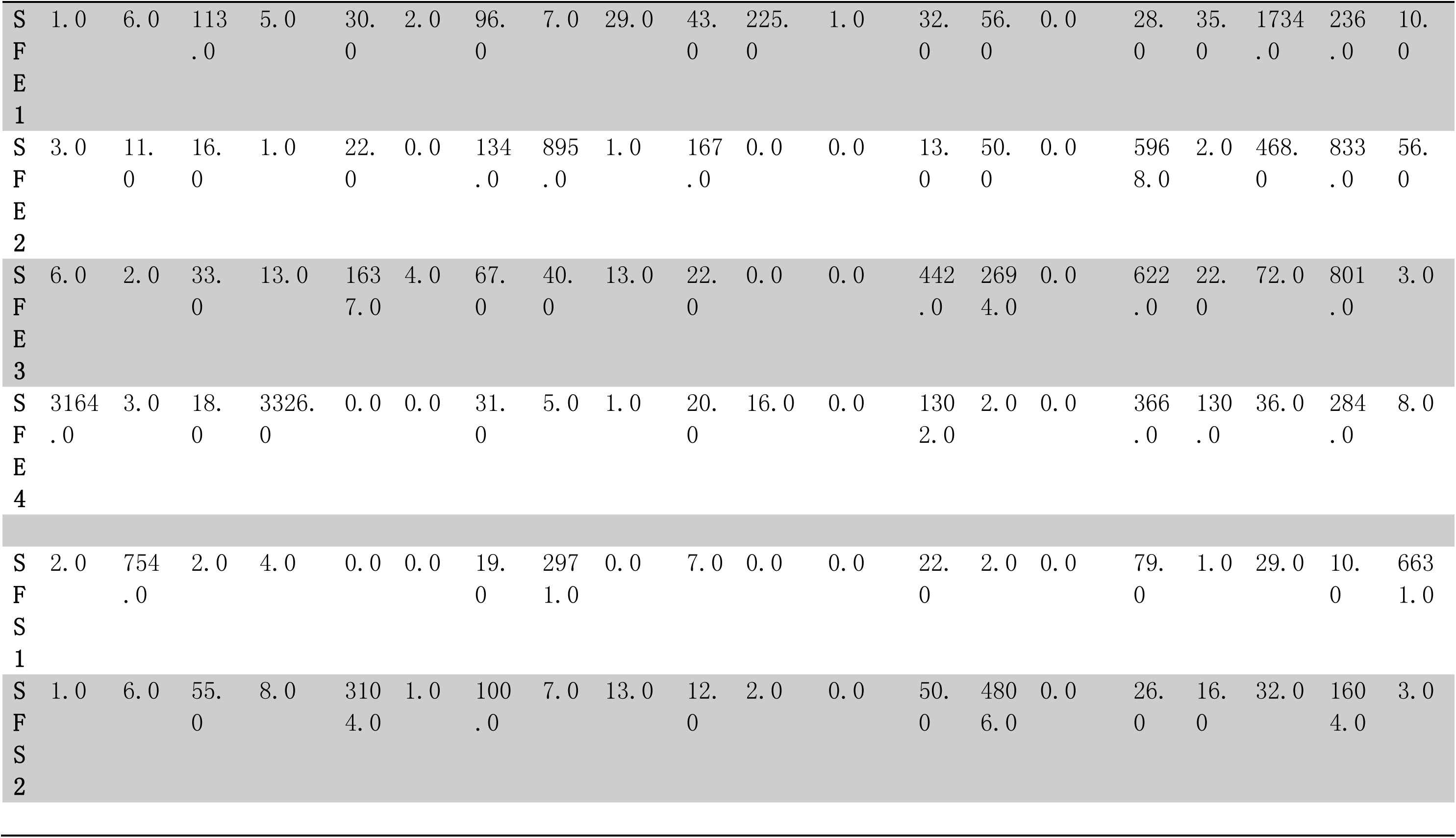
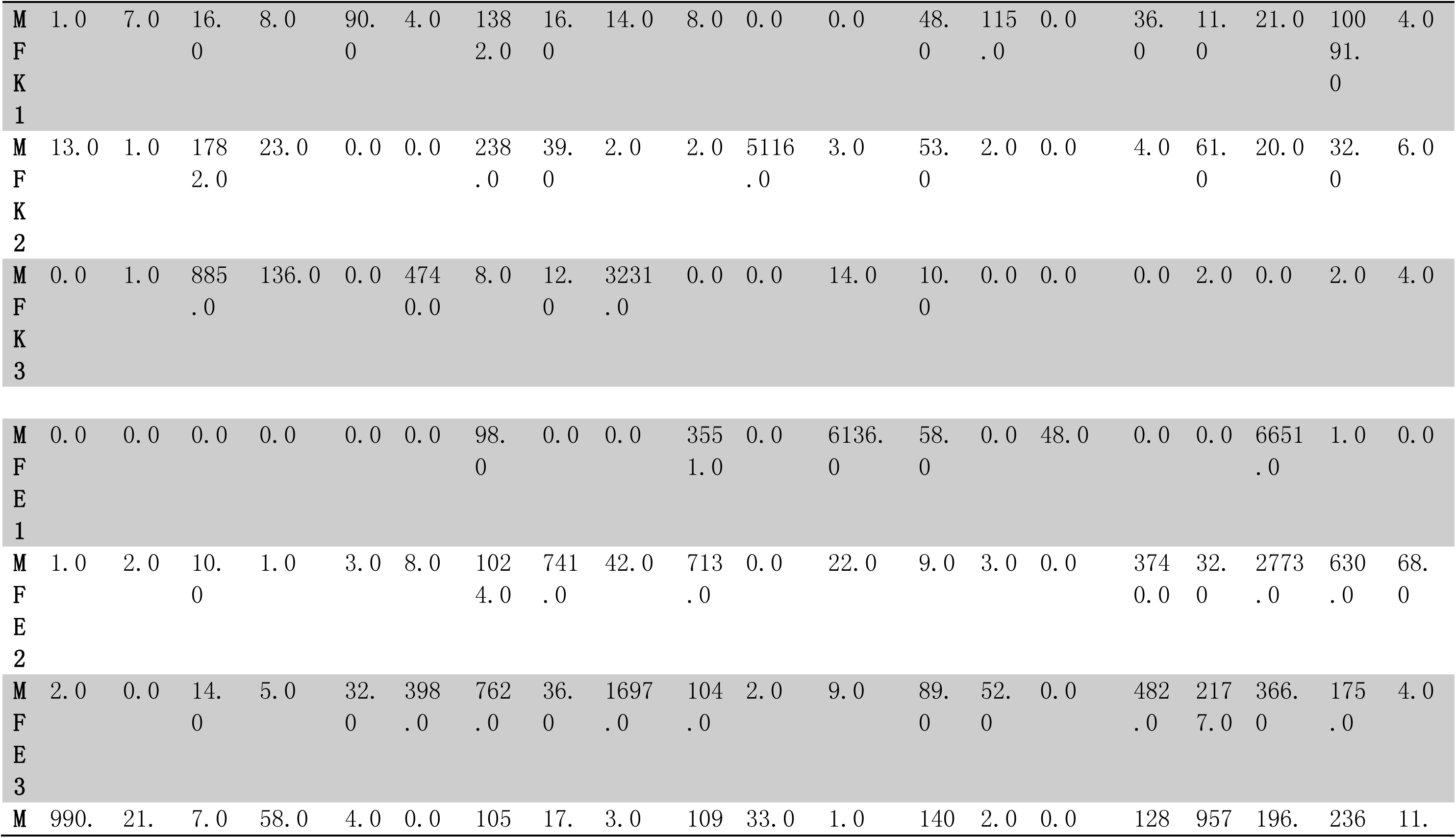
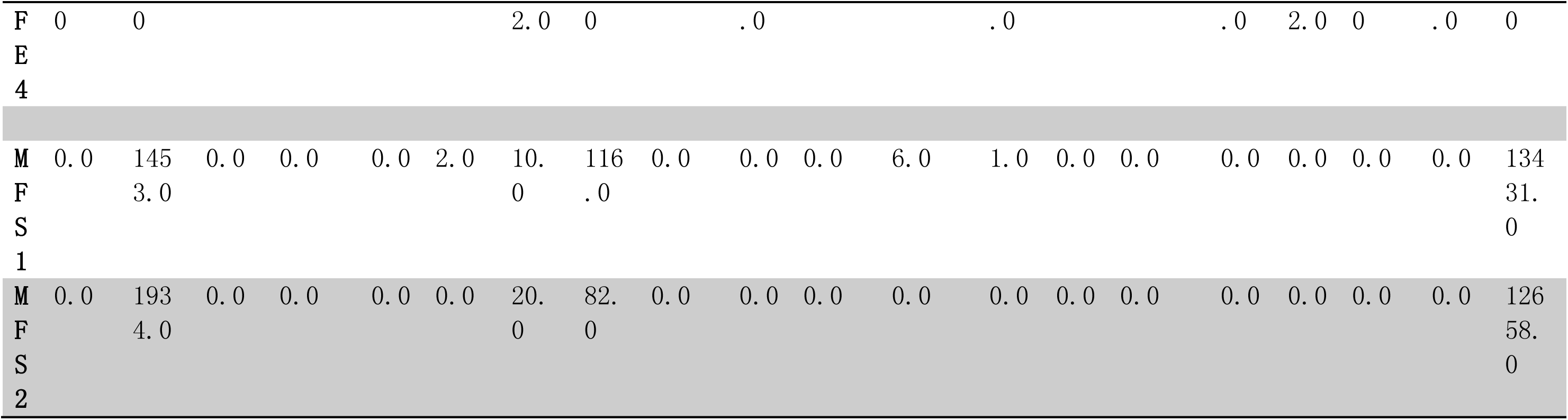

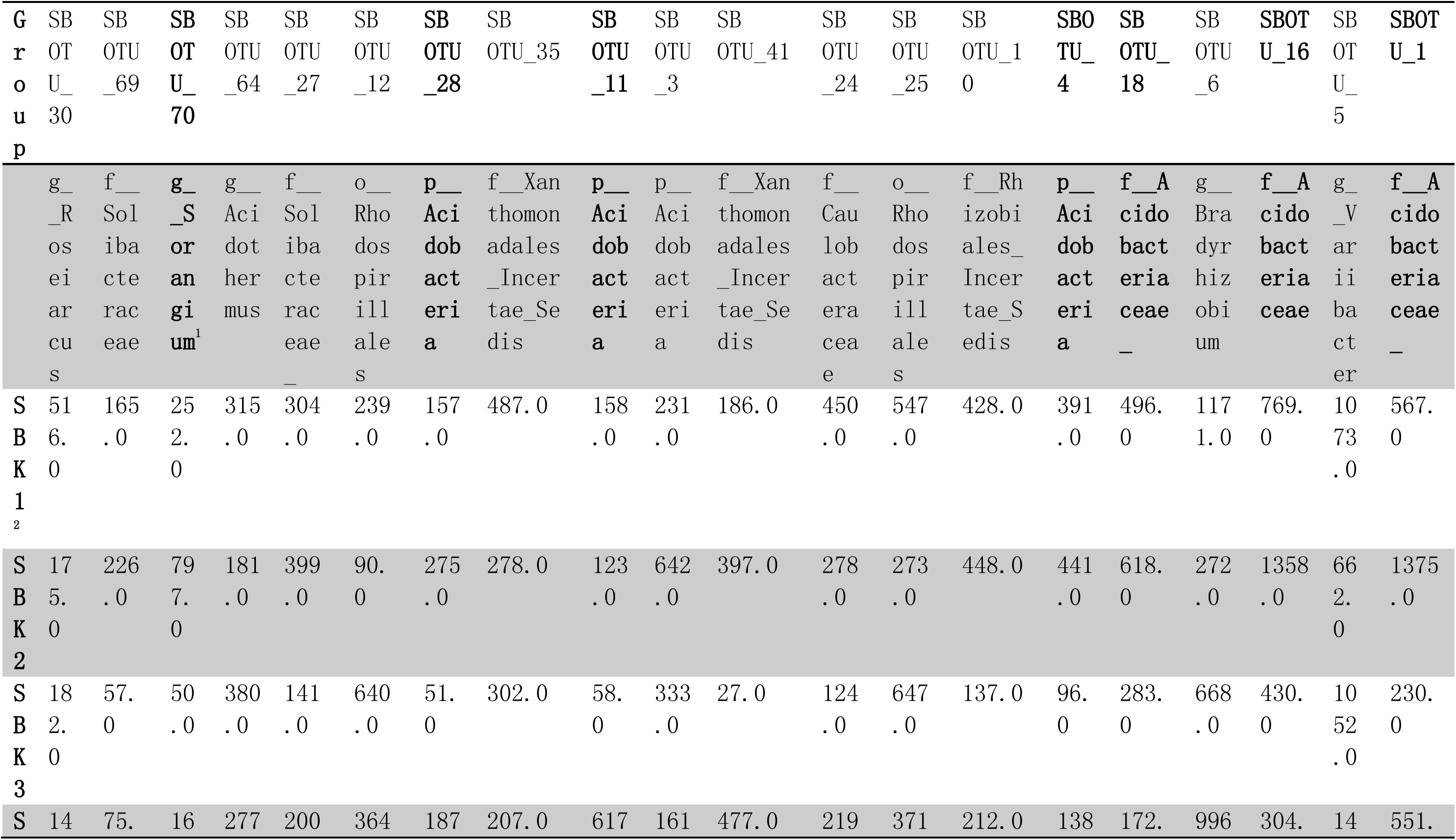
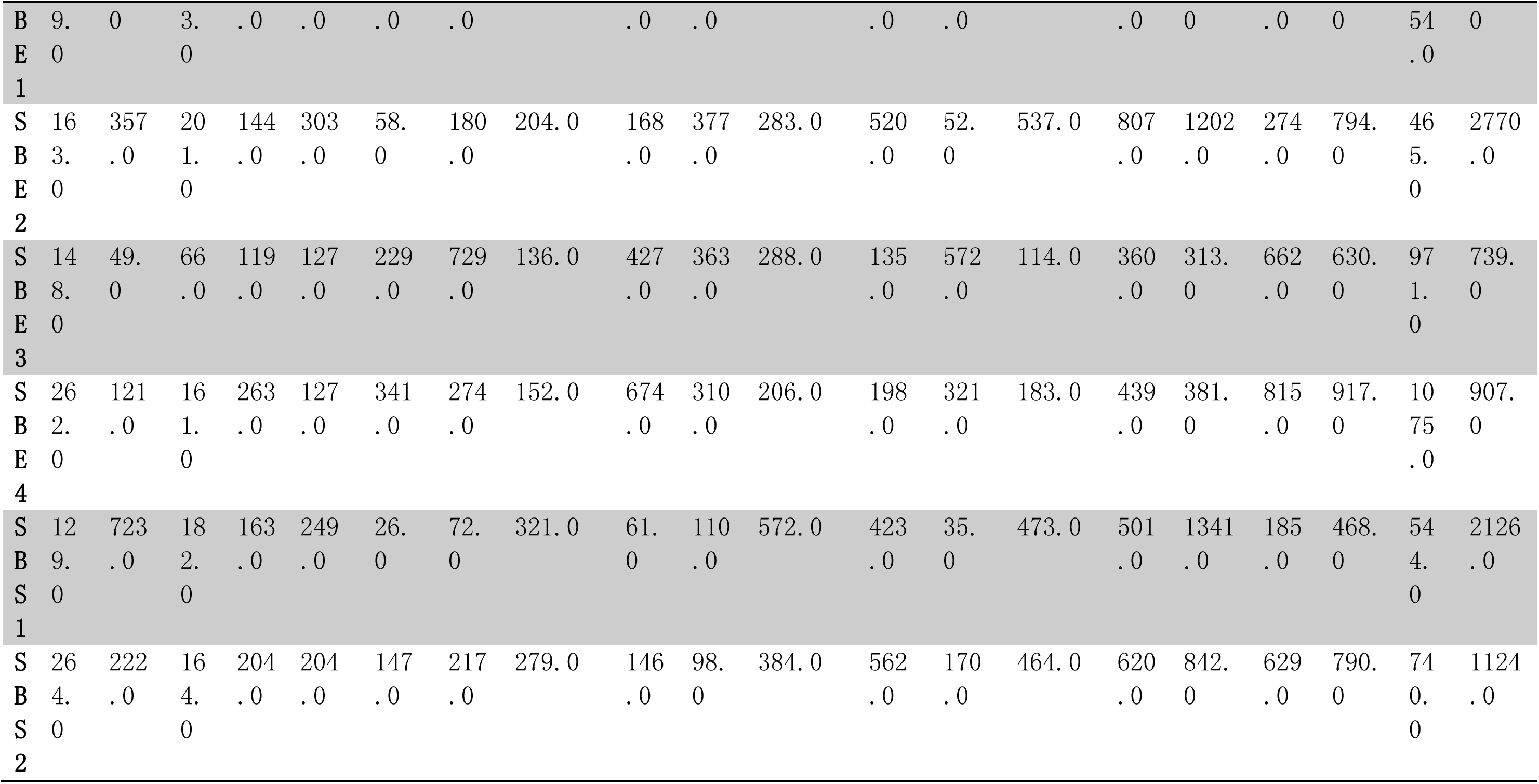

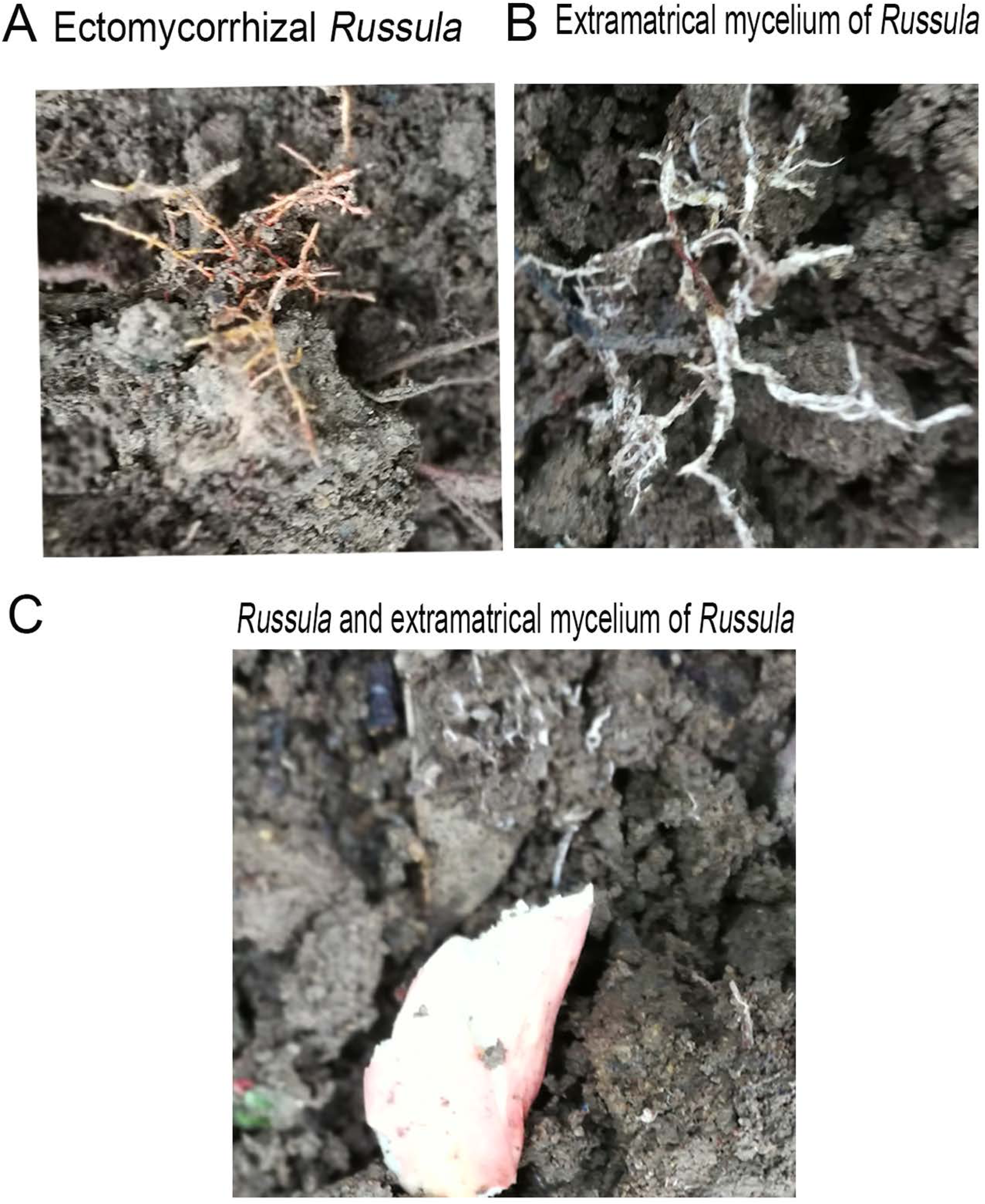

